# Notch-driven asymmetry dictates hair-cell behavior via a fate-specific kinase

**DOI:** 10.64898/2026.02.25.708020

**Authors:** Emily Atlas, Caleb C. Reagor, Brian Frost, Sapna Krishnakumar, A. J. Hudspeth, Adrian Jacobo

## Abstract

Cell-cell signaling and cell-fate decisions are essential for organ assembly, but how these molecular events translate into the physical properties and cell behaviors that drive development remains poorly understood. In the zebrafish lateral line, developing pairs of sensory hair cells undergo symmetry-breaking mediated by Notch signaling, leading to one cell becoming Notch-ON (receiver state) and its sibling becoming Notch-OFF (sender state). These cells then undergo coordinated movements that ensure that Notch-OFF cells are always positioned anterior to their Notch-ON sisters, and the Notch state of each sister ultimately determines the direction of hair bundle polarization. However, the cellular mechanisms and molecular programs that guide polarity-specific behaviors remain largely unknown. Here, using time-lapse imaging and a new 3D segmentation and tracking pipeline, we demonstrate that sister cells actively move in opposite directions – Notch-OFF cells migrate anteriorly while Notch-ON cells move posteriorly – enabling robust rotations when cells are initially mispositioned. Using single-cell RNA sequencing, we identify fate-dependent transcriptional programs and show that the differentially expressed kinase *stk32a* is required for proper navigation and hair bundle placement in Notch-ON cells. Strikingly, loss of *stk32a* reveals an underlying chiral bias in cell-pair rotations, suggesting the existence of an additional, previously unrecognized axis of symmetry breaking beyond the Notch-mediated fate decision.

## Introduction

Recent applications of high-resolution genomic sequencing technologies have helped reveal the intricate molecular networks that specify cell fate, but less is understood about how these networks send, receive, and process information to allow for collective cellular behaviors that coordinate tissue- and organ-level development (1, 2). Organogenesis requires not only the generation of the correct cell types but also the specification of cell numbers, cell-type ratios, and proper spatial organization. To understand the emergent mechanisms underlying development, homeostasis, and disease, it is crucial to understand the rules and constraints governing this specification. These developmental insights can ultimately reveal how novel tissue architectures evolve (3).

The difficulty of observing key morphogenetic events *in vivo* has limited our understanding of the complex, mechanochemical feedback mechanisms governing tissue development. The zebrafish lateral-line system can overcome many of these challenges, as it consists of relatively few cell types, is optically accessible over long periods of time, and is well-suited for dissecting cell behaviors (4– 7). The lateral line is composed of sensory organs called neuromasts that detect water flows across multiple axes, including the anteroposterior (AP) and dorsoventral (DV) axes (4, 8). Each neuromast contains sensory hair cells, each bearing on its apical surface a mechanosensory apparatus called the hair bundle. Hair cells come in two types, classified according to the direction of water flow they detect: in AP-oriented neuromasts, some sense flow from head-to-tail (caudad-polarized) while others sense flow from tail-to-head (rostrad-polarized) (8, 9).

During development, fate-restricted progenitor cells in the neuromast divide symmetrically to form pairs of daughter hair cells. Shortly after cytokinesis, sister hair cells form extensive contacts, which are actively maintained during early maturation (10–12). During this period, Notch1amediated lateral inhibition establishes opposite identities in sister cells; one cell expresses high levels of Delta ligand (sender state) and low levels of the Notch transcriptional effector, Notch Intracellular Domain (NICD), in the nucleus. Conversely, its sister expresses low levels of Delta ligands (receiver state) and has high levels of NICD (13). Hereafter, we will refer to these two alternative fates as Notch-OFF and Notch-ON, respectively.

This fate asymmetry determines the intracellular polarization of the hair bundle apparatus through differential expression of the transcription factor *emx2* (in the Notch-OFF cell) (13, 14), a factor that also regulates hair bundle polarization in the mammalian vestibular system (15–18). The Emx2+ (Notch-OFF) cell becomes caudad-polarized, and its sister (Notch-ON, Emx2-) becomes rostrad-polarized (13, 14). However, because of the stochastic nature of the lateral inhibition process (10, 13, 19), cell positioning is initially random; the more posterior cell of the pair acquires the Notch-OFF state about half of the time. When this happens, sister cells undergo a characteristic rotation around one another, swapping positions across the anteroposterior axis defined by core planar cell polarity signaling (8, 10, 13, 20). The combination of these two processes — stochastic lateral inhibition followed by coordinated cell-pair rotations — leads to a final configuration of the organ where Notch-OFF, Emx2+ cells locate to the anterior side of the organ, while their Notch-ON, Emx2-siblings locate to the posterior end.

Despite the clear role of Notch signaling in establishing the unique molecular identities between sister hair cells, it remains unresolved how this fate asymmetry is translated into precise cell-pair rotations. Previous studies have disagreed regarding the extent to which Notch signaling regulates these early cell behaviors based on analyses of *notch1a* mutants (10, 21). Furthermore, the downstream hair bundle polarity-effector *emx2* is not required for proper relative cell positioning, and its perturbation has a minimal impact on cell-pair rotation (20, 21). Therefore, it is unknown whether and how Notch-dependent molecular asymmetry impacts single-cell and cell-pair mechanical properties and behaviors during the early stages of zebrafish lateral-line hair cell development.

In this study, we interrogate Notch1a-mediated fate and its relationship to polarity-specific behaviors during neuromast assembly. By combining a 3D image segmentation pipeline with high-resolution single-cell RNA sequencing, we dissect the specific transcriptional programs driving hair cell behaviors. Our sequencing strategy enables us to overcome prior limitations and to resolve polarity-specific transcriptional programs downstream of Notch1a. We identify the kinase Stk32a as a requisite downstream effector of this pathway, demonstrating its role connecting fate asymmetry to polarized bundle formation and directed cell-pair rotation.

## Results

### Cell pair movement aligns sister cell centroids in three dimensions

To better visualize and analyze cell-pair rotational behaviors, we developed a semi-automated 3D segmentation and tracking pipeline (Figure 1A,B; Methods) that enables visualization of cell-pair rotations in 3D (Figure 1C,D; Supplemental Movies 1 and 2). Using 3D centroids extracted from our datasets, we measured the angle *θ* between the cell pair and the AP axis (Figure 1E, top). We observed three characteristic types of angular trajectories over our ten-hour datasets: (i) cell-pair swaps of ∼180°(Figure 1F), (ii) non-swapping pairs (Figure 1G), and (iii) off-axis divisions that generally realigned, and could either swap or retain the relative positioning of sisters (Figure 1H) (Supplemental Movies 3-5). These behaviors closely mirror cell-pair trajectories analyzed in two dimensions (10, 20–23), confirming that our 3D tracking pipeline robustly captures the same core rotational dynamics. However, unlike many 2D implementations, our pipeline does not require manual slice selection at each time point and can scale to analyze more experiments and behaviors than in previous studies.

**Fig. 1.**
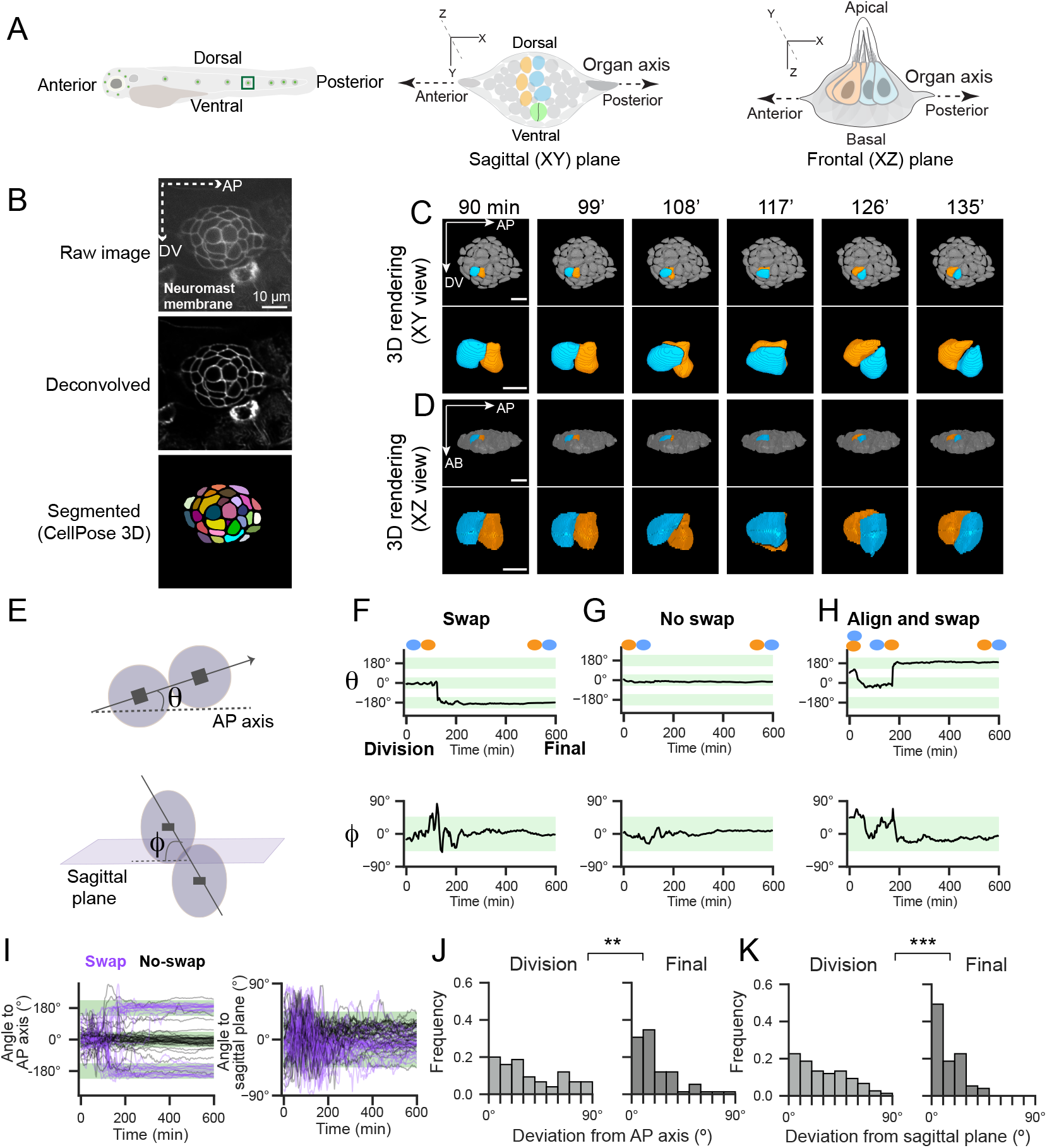
Semi-automated 3D analysis reveals hair-cell pair alignment dynamics. **(A)** Imaging geometry for time-lapse movies of hair cell development. *Left* : fish are mounted laterally, allowing imaging of neuromasts (example, boxed region). *Middle*: The imaging (sagittal) plane views the neuromast top-down, defining the Anteroposterior (AP) and Dorsoventral (DV) axes. *Right* : The frontal plane (XZ) reveals the apicobasal axis and depth of the organ. Hair cells are shown in orange and blue (newly developing pair in green); surrounding cells are in gray. **(B)** 3D image segmentation pipeline. A single sagittal section of a neuromast marked with membrane label *Tg(cldnb:lyn-mScarlet)* shows the progression from raw input (*top*) to deconvolved image (*middle*) to instance segmentation (*bottom*). Scale bar, 10 *µ*m. **(C)** Top-down and **(D)** side-view 3D renderings of neuromast during cell-pair rotation; times indicate minutes post cell division. Full neuromast in gray with cell pair in orange and blue (*top*); scale bar, 10 *µ*m. Zoomed in view of cell-pair (bottom); scale bar, 5 *µ*m. **(E)** Angle *θ* of cell pair with respect to AP axis and elevation angle *φ* of cell pair relative to the sagittal plane. **(F)** Example angular traces for *θ* (*top*) and *φ* (*bottom*) of a cell pair that swaps positions, **(G)** does not swap, and **(H)** divides off axis, aligns and swaps. Schematic of cell-pair positioning shown above each trace; light-green shading indicates *±*45° around AP axis (*top* panels) or sagittal plane (*bottom* panels). **(I)** Cell-pair angular traces with respect to AP axis (left) and sagittal plane (right) for 69 of 75 wild-type (WT) pairs; rare trajectories with angle to AP axis exceeding 320°were excluded for clarity. **(J)** Angular deviation of 75 WT cell pairs from AP axis and **(K)** from sagittal plane at division (initial) versus 10 hours post-division (final). Initial and final distributions compared using KS test (** p < 0.01, *** p < 0.001).

Beyond recapitulating known behaviors, we could also quantify aspects of cell-pair motion that were previously inaccessible. We examined the elevation angle *φ* of the cell-pair vector relative to the sagittal (XY) plane, a measure of how much the cell pair tilts out of the plane into the apicobasal dimension (Figure 1E, bottom). During the early stages of cell-pair development, large angular changes were observed, indicating that non-planar movements are common (Figure 1F-H). Notably, these deviations often coincided with periods of angular change along the AP axis. These out-of-plane movements can confound in-plane swaps when trajectories are viewed only in the sagittal projection. We therefore defined swaps in 3D by comparing sisters’ relative AP positions from initial alignment to the end of the time-lapse; pairs that reversed their AP order were classified as swaps (Methods, Figure S1A). In our dataset, 48% of 75 WT pairs underwent cell-pair swaps, close to the 50-60% previously reported (10, 20– 23).

In addition to undergoing targeted swaps, cell pairs also aligned their centroids in 3D over time. Pairs divided with a wide range of initial angles relative to the AP axis, but they ultimately settled along the axis at 0° (non-swapping) or ±180°(swapping in either the clockwise (positive) or counterclockwise (negative) direction) (Figure 1I, left). Cell-pair angles relative to the sagittal plane were also variable early but stabilized over time (Figure 1I, right). At the population level, sister-cell movements corrected initial misalignments both along the AP axis (Figure 1J) and with respect to the sagittal plane (Figure 1K), and this correction was observed in both swapping and non-swapping pairs (Figure S1B-C). Thus, cell-pair behaviors tend to enforce 3D alignment while also establishing a specific, final configuration through targeted cell-pair swaps.

### Notch state determines the direction of cell movement along the AP axis

To revisit the role of Notch asymmetry in the dynamics of this coupled cell behavior (10, 21), we generated a new dataset of cell-pair rotations in *notch1a*^*hzm17/hzm17*^ mutants. Neuromasts of these fish largely lack Notch-ON cells. In parallel, we generated a hair-cell-specific Notch gain-of-function line which mostly lacks Notch-OFF cells (*Tg(myo6b:NICD-P2A-mCherry)*, referred to as *Tg(myo6b:NICD))*. Although we did not observe mCherry expression in hair cells, mirroring expression issues reported previously for a similar transgene (20), transgenic larvae had a strong rostrad-polarity defect, indicating that expression of Notch1a Intracellular Domain (NICD) prior to the 2A tag was functional (13) (Methods). *Notch1a*^*hzm17/hzm17*^ mutants, as previously reported (14), have a strong caudad-polarity bias.

Cell pairs containing two Notch-OFF or two Notch-ON cells underwent swaps significantly less often than half of the time: 14% and 6.1% of the time, respectively (Figure 2A, Figure S2A,B). Notch-ON pairs exhibited an additional defect: they were unable to correct misalignments with respect to the AP axis, and cell pairs were significantly misaligned 10 hours post-division (Figure 2B). However, Notch-OFF and Notch-ON pairs were able to align to the sagittal plane over time (Figure S2C), suggesting that Notch signaling specifically impacts planar movements along the AP axis.

**Fig. 2.**
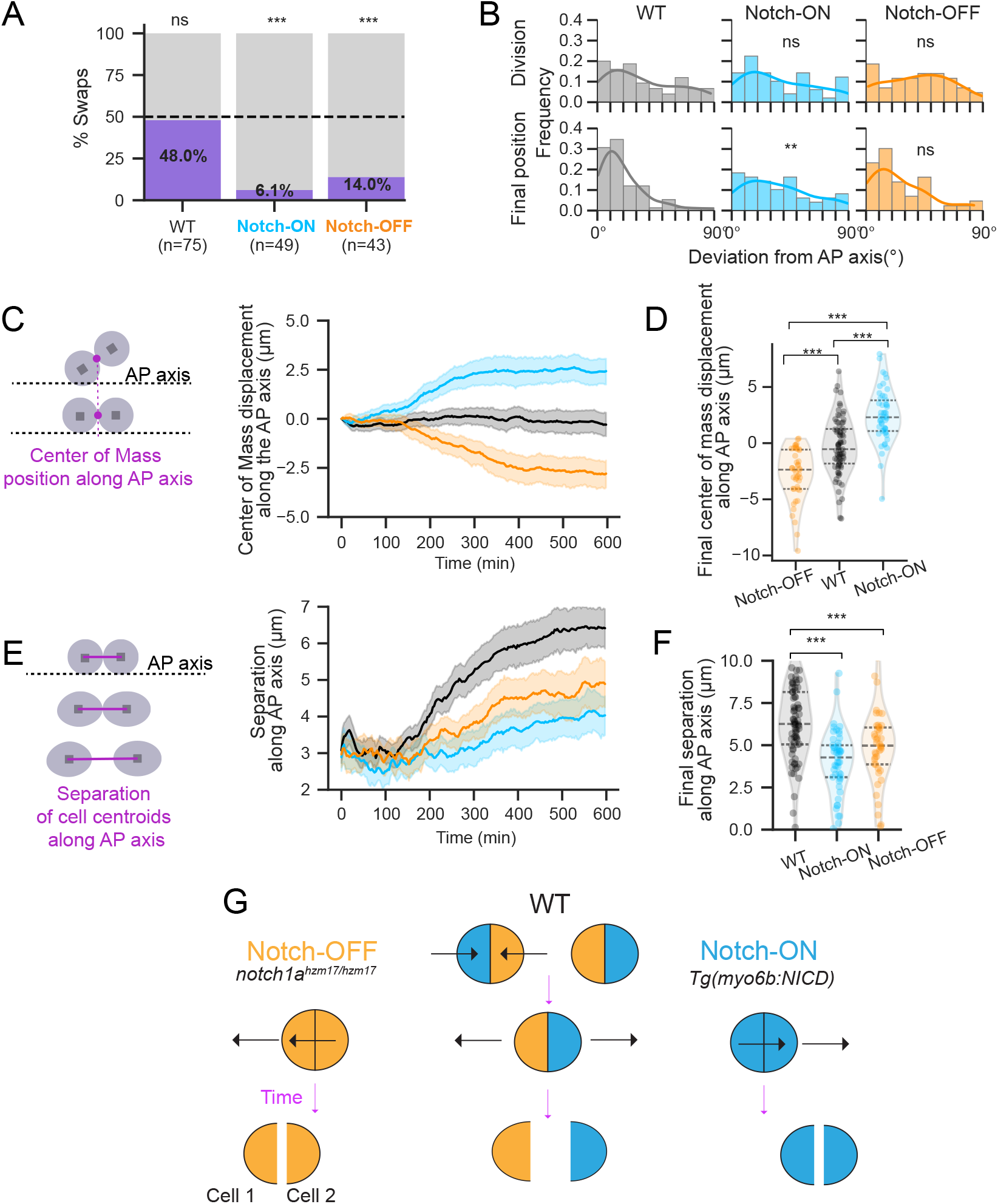
Notch1a-mediated symmetry breaking is required for robust cell-pair rotations. **(A)** Percentage of cell-pair swaps. Swapping occurs in roughly half of WT pairs (48.0%) but is significantly reduced in Notch-ON (6.1%) and Notch-OFF (14.0%) conditions. Dashed line indicates random chance (50%). (WT, n=75 pairs from 44 neuromasts across 20 fish, Notch-ON, n=49 pairs from 29 neuromasts, 16 fish, and Notch-OFF, n=43 pairs from 31 neuromasts, 18 fish). Each condition was compared to a 50% swap probability (binomial test); ***p < 0.001, ns p ≥ 0.05. WT data plotted from Figure 1 for comparison in this and subsequent panels. **(B)** Angular deviation of cell pairs from AP axis at division (*top*) and final (*bottom*). Distributions compared to WT at initial and final timepoints (KS test); ** p < 0.01, ns p ≥ 0.05, Holm-corrected. **(C)** Center of mass displacement of cell pair along the AP axis over time. Mean curves of all cell pairs are shown; error bands represent *±*95% confidence intervals. **(D)** Final center of mass displacement along the AP axis; *** p < 0.001 (one-way ANOVA with Tukey’s HSD correction). **(E)** Separation between sister cell centroids along the AP axis over time. Mean curves with *±*95% confidence intervals are shown. **(F)** Final separation between sister cell centroids; ***, p < 0.001 (Mann-Whitney U test, Holm-corrected). Schematic model of predicted movements in cell pairs. WT pairs (middle) consist of one Notch-OFF (orange) and one Notch-ON (blue) cell; opposing directional biases promote either cell-centroid separation or swapping depending on initial configuration. Pairs of two Notch-OFF (left) or two Notch-ON cells (right) drift as a unit and exhibit delayed separation.

The directionality of cell-pair movements was also perturbed. Whereas WT cell pairs tended to maintain the location of their center of mass at division over time without drifting along the AP axis, the center of mass of Notch-OFF pairs tended to drift anteriorly, as previously described using another *notch1a* mutant allele (10) (Figure 2C). Strikingly, Notch-ON pairs drifted posteriorly, opposite to the observed anterior drift of Notch-OFF pairs (Figure 2C,D). This fate-dependent drift was accompanied by disrupted sister-cell separation along the AP axis (Figure 2E,F). At the single-cell level, symmetric Notch-OFF or Notch-ON pairs consist of two sisters that move in the same direction, whereas WT pairs contain one Notch-ON and one Notch-OFF cell that move in opposing directions (Figure S2D-G, Figure 2G).

Together, these results suggest that opposing, Notch-driven, polarity-specific movements of individual sisters underlie robust pair alignment, rotation, and separation. Moreover, the strong impact of expression of the transcriptional effector NICD on cell behavior suggests that Notch1a sets the direction of movement through downstream transcriptional changes within the hair cell itself.

### Notch signaling defines distinct transcriptional trajectories that underlie polarity-specific hair-cell fates

We next sought to identify the transcriptional changes associated with Notch1a signaling in hair cells. Although numerous single-cell RNA sequencing (scRNA-seq) studies have been performed on the zebrafish lateral line (14, 20, 24, 25), these analyses have failed to identify polarity-specific trajectories. Therefore, we performed scRNA-seq on fluorescently sorted hair cells (Figure S3) from *notch1a*^*hzm17/hzm17*^ mutants, their heterozygous and WT siblings, and other WT animals (Figure 3A). Following quality control and selection of lateral line hair cells (Methods, Figures S4, S5), we examined the expression of the polarity-specific transcription factor *emx2*. Despite robust detection, *emx2*-positive and *emx2*-negative cells did not form separate clusters on the UMAP; instead, hair cells grouped by maturity and genetic background (Figure S5E,F). However, differential gene expression analysis with DESeq2 (26–28) identified Notch-dependent transcriptional changes (Figure S6A-B, Supplemental Tables 1 and 2), including the downregulation of canonical Notch signaling targets (29, 30) (*e*.*g. hey2* and *her4*.*1*).

**Fig. 3.**
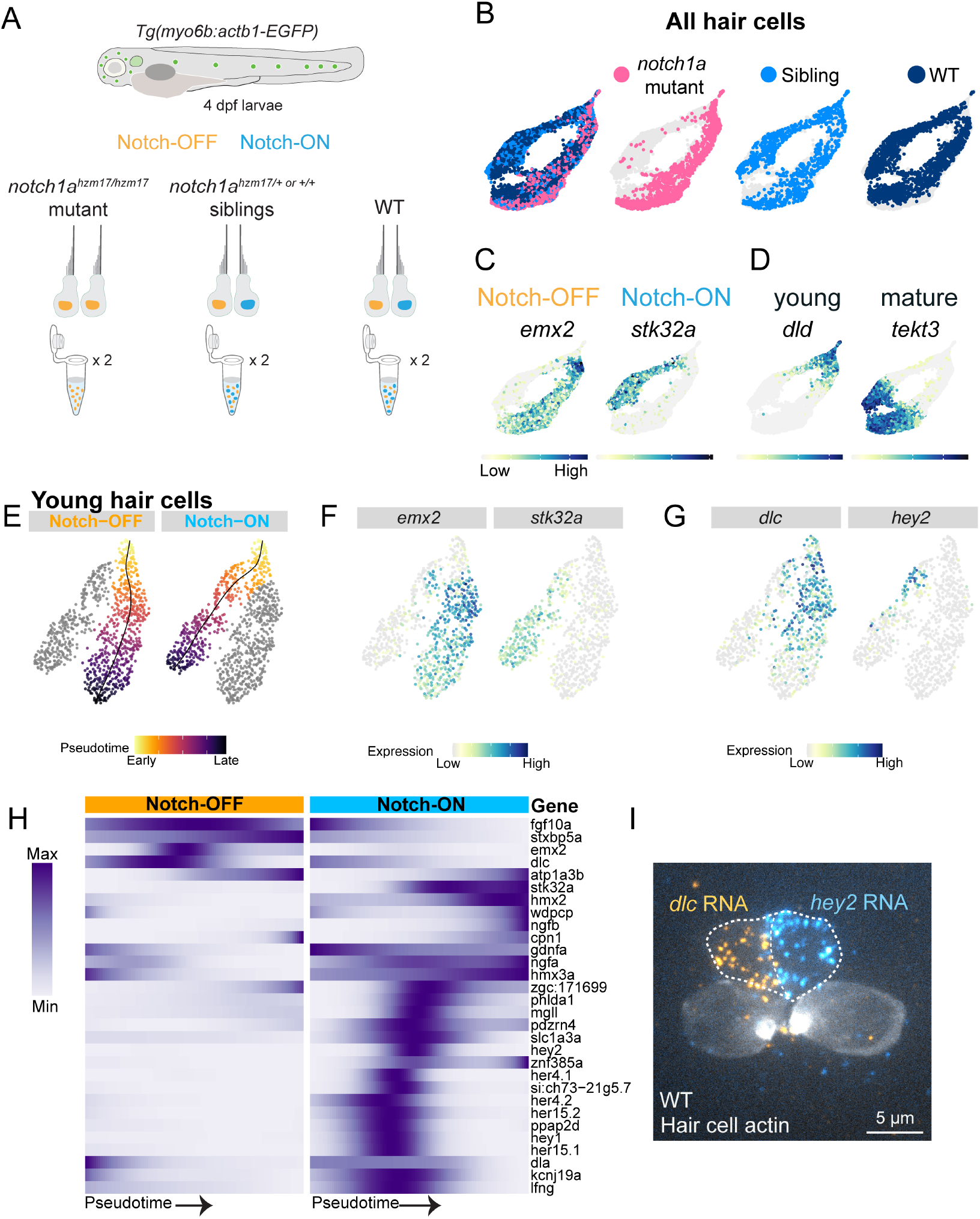
Notch signaling defines distinct transcriptional trajectories that underlie polarity-specific hair-cell states. **(A)** scRNA-sequencing workflow. Hair cells were isolated from WT and *notch1a*^*hzm17/hzm17*^ mutant fish and their siblings. While heterozygous siblings and WT fish have both caudad- and rostrad-polarized cells, mutants predominantly have caudad-polarized hair cells. **(B)** UMAP visualization of all hair cells following a targeted dimensionality reduction for polarity-specific expression (see Methods and Figure S6). **(C)** Feature plots showing SCT-normalized expression of previously identified polarity-specific transcription factor *emx2* and the polarity-specific kinase *stk32a*, and **(D)** young and mature hair cell markers *dld* and *tekt3*, respectively. **(E)** Trajectory analysis of subsetted and reclustered young hair cells; UMAP overlaid with Slingshot-determined Notch-OFF and Notch-ON lineages (black line) and pseudotime values (colormap). **(F-G)** Expression patterns of example lineage-specific genes. Expression patterns (min-max normalized) of top 30 lineage-specific genes identified by TradeSeq (see Methods). **(I)** *dlc* (orange) and *hey2* (cyan) RNA visualized through HCR *in situ* hybridization. Maximum-intensity projection of a 2-dpf neuromast is shown contrast-adjusted to visualize expression patterns, with nascent hair-cell pair outlined by a dotted line and identified by *Tg(myo6b:actb1-EGFP)* expression (grayscale). Scale bar, 5 *µ*m.

To find a polarity-specific signature in the single-cell data, we applied a targeted dimensionality-reduction approach (Methods). In brief, we performed principal component analysis and determined which PC scores covaried with the polarity-specific transcription factor *emx2* (Figure S6C-D). These PCs were used to select genes with the highest loadings, which we then used to generate a new polarity-specific UMAP embedding. This approach revealed two distinct trajectories in the UMAP embedding, with most *notch1a*-mutant cells in one branch, whereas sibling and WT cells were distributed across both branches (Figure 3B). The generated UMAP shows a separation of young and mature hair cells (Figure 3D), and *emx2*+ hair cells largely separate onto the putative Notch-OFF branch (Figure 3C). Recent work in the mouse ear demonstrated that the kinase *stk32a* is differentially expressed in the vestibular system, specifically in hair cells lacking *emx2* (31). *stk32a* showed strong differential expression in our dataset as well (Supplemental Table 1), suggesting it is polarity-specific in the zebrafish as well. Therefore, we visualized *stk32a* expression in our embedding and indeed observed its specific expression on the Notch-ON/*emx2-* negative branch (Figure 3C).

Since cell-pair rotations occur in young hair-cell pairs, we next subset and recomputed a polarity-specific embedding for young hair cells only. We then performed trajectory analysis, revealing a putative branching point in very young hair cells (Figure 3E-G), and compared lineage-specific expression patterns as young hair cells matured (Supplemental Table 3). We visualized the dynamics of the top polarity-specific genes acting downstream of Notch on a heat map (Figure 3H) and observed a transient activation of canonical Notch target gene expression specifically along the Notch-ON branch. To relate our pseudotime analysis to the timing of expression in young hair cells, we used *in situ* hybridization chain reaction (HCR) to simultaneously visualize the expression of *hey2* – a canonical Notch-signaling target that was significantly downregulated in the *notch1a* mutant – and *dlc*, a delta ligand that was upregulated. As expected, we observed polarity-specific expression of both genes in the opposite sister cells of an immature WT pair (Figure 3I), indicating that our pseudotime analysis identified genes expressed around the same time as cell-pair rotations.

### Stk32a mediates polarity-specific hair-bundle alignment downstream of Notch

To understand whether any of the identified genes could impact cell polarization during hair cell development, we then performed a small-scale candidate screen using an F0 crispant approach (see Methods) (32, 33), prioritizing non-transcription factors to find polarity effectors rather than other components of the gene regulatory network. We analyzed hair-bundle orientation in young neuromasts as our primary readout for polarity defects. Screening for bundle misorientation allowed us to quickly assess phenotypes caused by defects in cell polarization and upstream cell-pair rotations. Of the genes we tested (Supplemental Table 4), only knockdown of the serine-threonine kinase *stk32a* produced robust defects in hair-bundle polarity without disrupting neuromast deposition, making it a promising candidate for further analysis of its role in hair cell polarization.

We used CRISPR/Cas9 to generate a stable *stk32a*^*ru800/ru800*^ mutant line (see Methods). In young, 2 dpf, homozygous *stk32a*^*ru800/ru800*^ animals, we observed a clear caudad-polarity bias, along with a substantial fraction of hair cells whose bundles failed to align along the anteroposterior axis (Figure 4A-B). Notably, heterozygous *stk32a*^*ru800/+*^ siblings rarely possessed such off-axis hair bundles. Furthermore, *notch1a*^*hzm17/hzm17*^ mutants, which also displayed a strong caudad-polarity bias (Figure 4C), remained aligned along the AP axis. Therefore, *stk32a* loss-of-function only partially phenocopies the effects of *notch1a* disruption. We also generated a *Tg(myo6b:stk32a-EGFP)* line to express *stk32a* in all hair cells. At 2 dpf, these neuromasts had an abundance of rostrad-polarized cells in addition to cells whose hair bundles were not aligned to the AP axis (Figure 4D-E). In contrast, *Tg(myo6b:NICD)* hair cells did not often polarize dorsally or ventrally (Figure 4F). In older, 5 dpf larvae, the off-axis polarity defects were decreased compared to 2 dpf larvae, but the caudad-bias in *stk32a* mutants and the rostrad-bias in *Tg(myo6b:stk32a)* larvae persisted (Figure S7). Thus, perturbing *stk32a* disrupts the balance of hair bundle-polarization, but not identically so to *notch1a* perturbation.

**Fig. 4.**
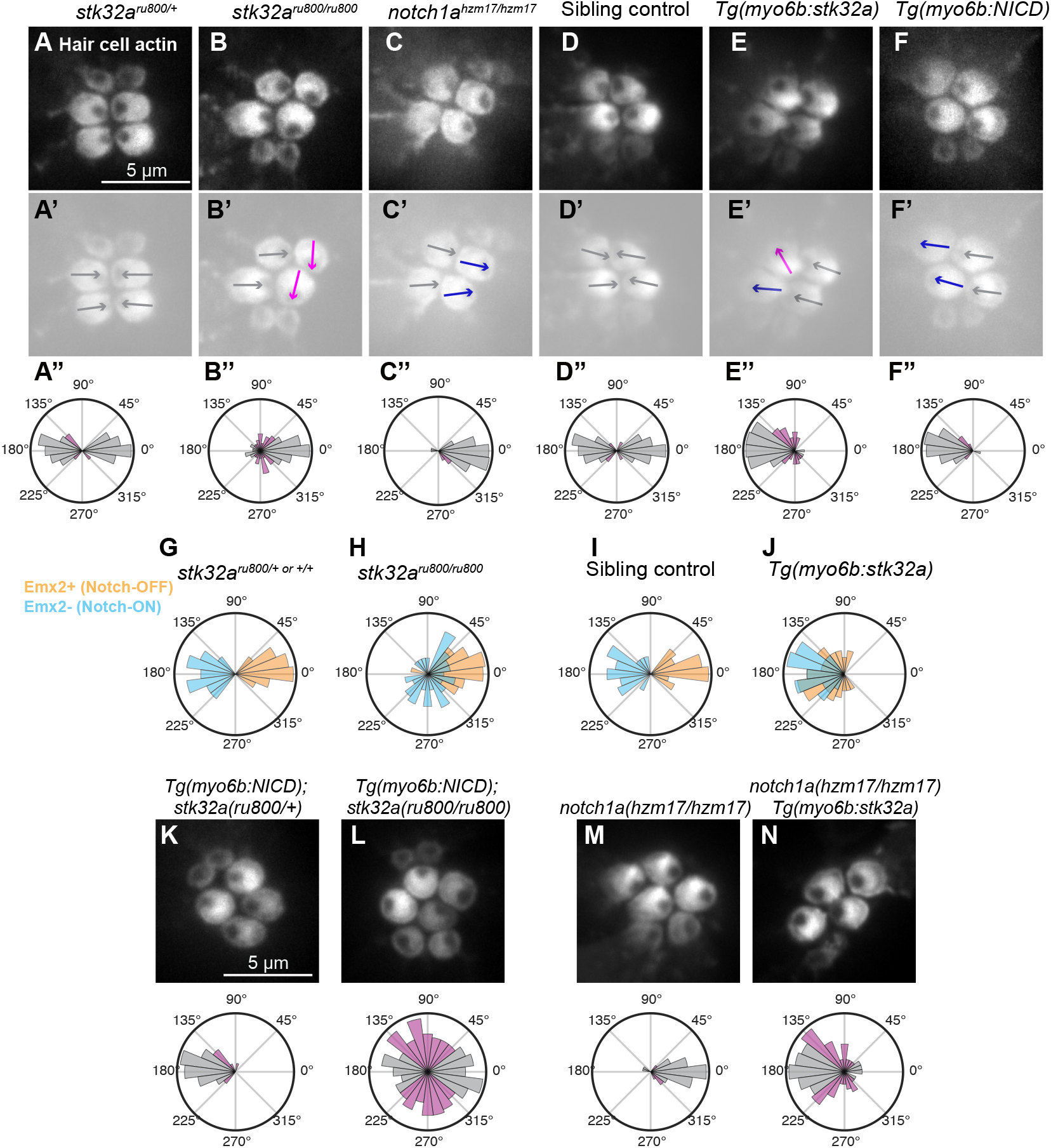
*stk32a* asymmetry is required for mirror-image hair bundle polarization downstream of Notch1a. **(A-F)** Top-down views of apical surface of neuromast hair cells labeled with *Tg(myo6b:actb1-EGFP)*, showing hair bundle polarities in 2-dpf larvae for the indicated genotypes. Scale bar, 5 *µ*m. (**A’–F’**) Same images, contrast adjusted and arrows overlaid indicating individual hair-bundle polarity. Gray arrows denote bundles aligned to the AP axis and in the correct direction (close to 0°or 180°depending on position); pink arrows denote off-axis bundles (close to 90°or 270°); navy blue arrows denote bundles aligned to the AP axis but in the reversed direction, breaking mirror-symmetry. **(A’ ‘-F’ ‘)** Rose plots of hair bundle polarization for each genotype; bar color indicates orientation relative to the AP axis (gray: AP-aligned; magenta: off-axis). (A) *stk32a*^*ru800/+*^ siblings (n = 106 hair cells from 27 neuromasts in 8 larvae). (B) *stk32a*^*ru800/ru800*^ (n = 155 hair cells from 39 neuromasts in 12 larvae). (C) *notch1a*^*hzm17/hzm17*^ mutants (n = 158 hair cells from 33 neuromasts in 10 larvae). (D) Sibling non-expressing controls (n = 130 hair cells from 32 neuromasts in 8 larvae). (E) *Tg(myo6b:stk32a)* (n = 148 hair cells from 39 neuromasts in 10 larvae). (F) *Tg(myo6b:NICD)* (n = 142 hair cells from 32 neuromasts in 8 larvae). **(G-J)** Hair-bundle polarity distributions in Emx2+ (Notch-OFF) and Emx2-(Notch-ON) hair cells in each genotype displayed. For each condition, a rose plot with Emx2+ and Emx2-cells is shown overlaid with Emx2+ cells in orange and Emx2-cells in blue. (G) *stk32a*^*ru800/+* or +/+^ siblings (n = 102 total hair cells from 22 neuromasts in 9 fish) (H) *stk32a*^*ru800/ru800*^ mutants (n = 91 total hair cells from 25 neuromasts in 10 fish) (I) Sibling non-expressing controls (n = 62 total hair cells from 20 neuromasts in 11 fish) (J) *Tg(myo6b:stk32a)* (n = 90 total hair cells from 24 neuromasts in 11 fish). **(K–N)** Apical surface of neuromast hair cells (*top*) and rose plots quantifying hair cell polarity *(bottom*) in 2-dpf larvae, shaded as in panels A-F. **(K)** *Tg(myo6b:NICD)*; *stk32a*^*ru800/+*^ (n = 140 hair cells from 35 neuromasts in 9 larvae). **(L)** *Tg(myo6b:NICD)*; *stk32a*^*ru800/ru800*^ (n = 162 hair cells from 43 neuromasts in 11 larvae). **(M)** *notch1a* ^*hzm17/hzm17*^ (n = 115 hair cells from 28 neuromasts in 8 larvae). **(N)** *notch1a* ^*hzm17/hzm17*^; *Tg(myo6b:stk32a)* (n = 139 hair cells from 40 neuromasts in 12 larvae). Scale bar, 5 *µ*m.

We next confirmed that Notch1a regulates *stk32a*. HCR *in situ* hybridization showed that *stk32a* RNA was expressed in all hair cells of *Tg(myo6b:NICD)* larvae but specifically in the rostrad-polarized population of WT neuromasts (Figure S8A-D). Furthermore, *stk32a* perturbation does not impact the prevalence of Emx2-positive hair cells, further indicating that Notch signaling works upstream of *stk32a* (Figure S8E-F).

Downstream of Notch, *stk32a* perturbation impacted the polarization of Notch-OFF and Notch-ON cell types differently. In control larvae, Notch-OFF cells (Emx2+) were caudad-polarized, and Notch-ON cells (Emx2-) were rostrad-polarized (Figure 4G). In *stk32a* mutants, on the other hand, while Notch-OFF cells remained caudad-polarized, Notch-ON cells showed strong misorientation, often aligning along the dorsoventral axis (Figure 4H). Conversely, in *Tg(myo6b:stk32a)* larvae, Notch-OFF cells were the ones strongly mispolarized (Figure 4 I,J). Together, these results suggest that *stk32a* is required for proper hair-bundle polarization in Notch-ON cells and is sufficient to mispolarize Notch-OFF cells, identifying Stk32a as a key driver of hair-bundle polarization in the zebrafish neuromast.

We next asked whether removing *stk32a* could alter the rostrad polarity-bias induced by transgenic NICD expression. In *Tg(myo6b:NICD*); *stk32a*^*ru800/+*^ larvae, in which both sisters adopt the Notch-ON state, hair bundles remained strongly rostrad-polarized (Figure 4K) like *Tg(myo6b:NICD*) larvae (Figure 4F). In contrast, *Tg(myo6b:NICD); stk32a*^*ru800/ru800*^ neuromasts showed a broad, randomized distribution of hair bundle polarization (Figure 4L). These results further demonstrate that *stk32a* is required for proper alignment of Notch-ON hair bundles along the organ’s axis and in the rostrad direction. We next tested whether hair-cell-specific *stk32a* expression alters polarity distributions in Notch-deficient hair cells. In *notch1a*^*hzm17/hzm17*^ mutants, *Tg(myo6b:stk32a)* expression shifted hair-bundle polarity away from a strong caudad bias (Figure 4M) toward a rostrad bias (Figure 4N). Unlike the randomized polarity observed upon *stk32a* removal in NICD-expressing hair cells, ectopic *stk32a* expression in Notch-OFF cells produced a strongly biased, albeit not tightly constrained, polarity distribution.

In the mouse utricle, polarity reversal through Stk32a works in parallel to cues provided by the core PCP pathway (31), raising the question of whether Stk32a serves a similar role in the zebrafish neuromast. To assess this, we examined the localization of the PCP protein Vangl2. In control neuromasts, Vangl2 localized to the posterior edge of hair cells, as previously reported (8, 13) (Figure S9A). In *Tg(myo6b:stk32a)* neuromasts, Vangl2 remained posteriorly enriched, including in cells with off-axis bundle orientations (Figure S9B-C). Thus, perturbing *stk32a* asymmetry does not disrupt PCP protein localization. Because *stk32a* is expressed specifically in Notch-ON cells, it enables cell-type-specific interpretation of PCP directional cues, allowing two sisters exposed to the same tissue-level polarity signal to respond in opposite ways.

### *stk32a* loss-of-function impacts cell-pair rotational behavior and uncovers a chiral rotation bias

*stk32a* is necessary for Notch-ON cells to polarize their hair bundles along the AP axis and sufficient to alter the polarization of Notch-OFF cells. We returned to our analysis of cell-pair rotations to determine if *stk32a* also impacts the ability of cells to move directionally downstream of Notch. As discussed in the introduction, in young WT neuromasts nearly all Emx2+ (Notch-OFF) cells are localized on the anterior side of the organ. In *stk32a* mutants, 73% of Emx2+ (Notch-OFF) hair cells were properly placed, a significant reduction, though not the 50% that would be expected if all rotation events had failed (Figure S10A,B). Hair-cell specific *stk32a* expression (*Tg(myo6b:stk32a))* on the other hand, did not have a significant effect on cell positioning (Figure S10B).

To confirm the effect of *stk32a* perturbation on cell-pair rotations, we conducted live imaging of *stk32a*^*ru800/ru800*^ mutant and *Tg(myo6b:stk32a)* pairs and compared their dynamics to WT. Mutant pairs swapped positions 30.5% of the time, a reduction from the expected 50% (Figure 5A), but a milder defect than that seen under *notch1a* perturbation (Figure 2A). *Tg(myo6b:stk32a)* pairs swapped 40% of the time (Figure 5A). Furthermore, *stk32a* mutant and *Tg(myo6b:stk32a)* pairs did not exhibit defects in AP axis alignment by the end of the time-lapse (Figure 5B). Despite modest defects in cell-pair angular behavior, we noticed that *stk32a* mutants exhibited a strong bias toward clockwise (CW) rotations when they swapped (Figure 5C), whereas WT and *Tg(myo6b:stk32a)* pairs showed no such directional preference. This chirality was surprising, so we next sought to understand how it could arise.

**Fig. 5.**
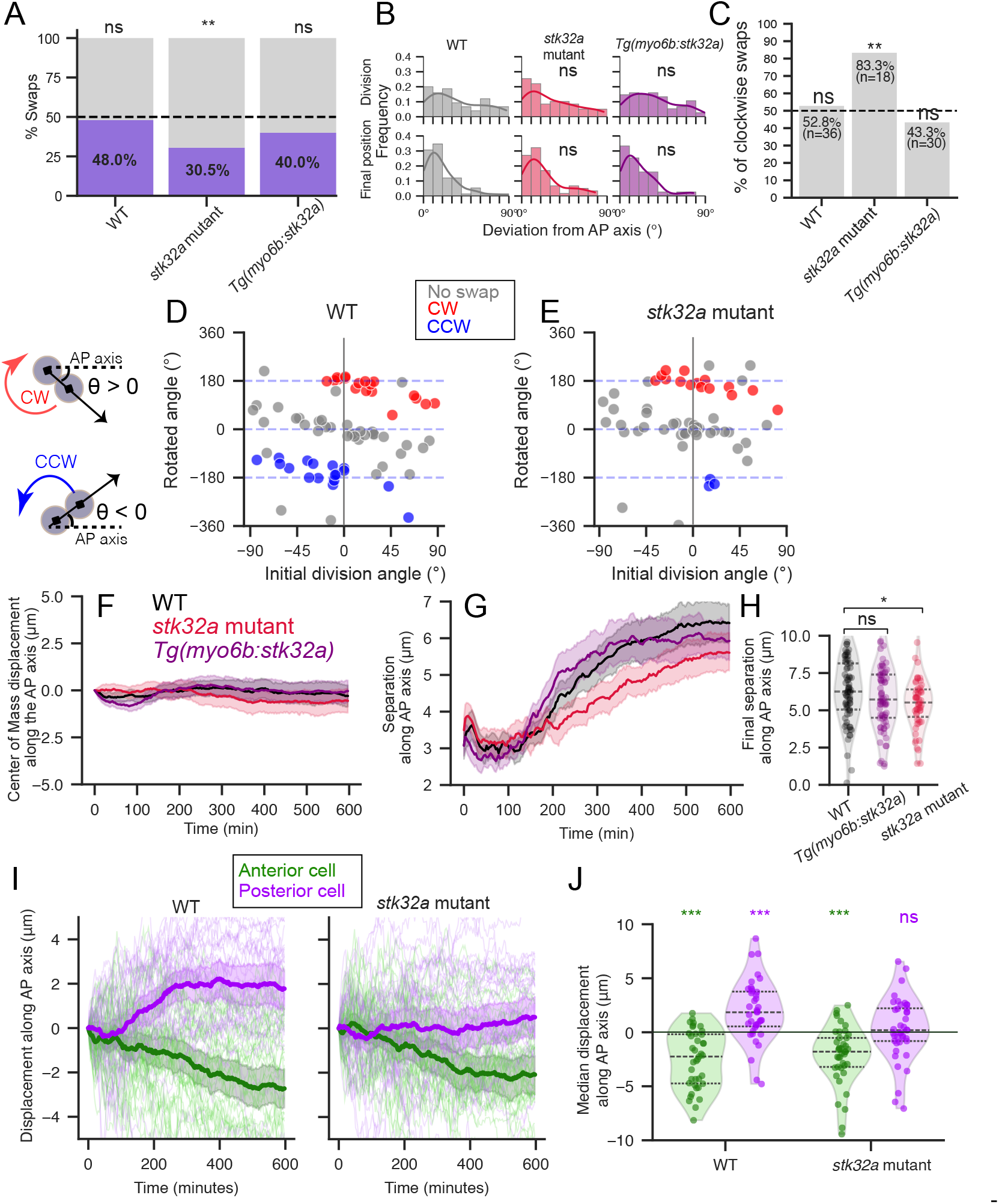
*stk32a* perturbation impacts the behavior of developing cell pairs. **(A)** Percentage of swaps in *stk32a* mutant (n = 59 pairs from 37 neuromasts across 19 fish) and *Tg(myo6b:stk32a)* (n = 75 pairs from 38 neuromasts across 18 fish). WT data are replotted from Figure 1 in this and subsequent panels for comparison. Each condition compared to a 50% swap probability using binomial test; **p < 0.01, ns p ≥ 0.05. **(B)** Deviation of pairs from AP axis at division and final timepoints, compared to WT at each timepoint; ns p ≥ 0.05 (KS test, Holm-corrected). **(C)** Percentage of swapping pairs that rotate clockwise (CW) compared to 50% CW by binomial test; ** p < 0.01, ns p ≥ 0.05. **(D)** Initial division angle plotted against net rotation angle in WT pairs; pairs colored by whether they undergo swaps (CW: red, CCW: blue, no swap: gray). Rare pairs that rotated ≥ 360°were excluded (3/75 pairs). **(E)** Same as (D) for *stk32a*-mutant cell pairs (1/59 pairs excluded). **(F)** Translation of the center of mass of the cell pair along the AP axis over time. **(G)** Distance between cell centroids of sister cells along the AP axis. For panels F and G, the mean and error bands representing the 95% CI are plotted. **(H)** Final cell pair separation along the AP axis; * p < 0.05, ns p ≥ 0.05 (Mann-Whitney U test, Holm-corrected). **(I)** For pairs that did not undergo swapping, displacement along the AP axis was plotted separately for the anterior-most and posterior-most cells, shown in green and purple, respectively. Traces are shown for WT (*left*) and *stk32a* mutants (*right*). Individual cell traces are shown with pale lines, and the mean trajectory is plotted, with 95% confidence intervals. **(J)** Median AP-axis displacement for anterior (green) and posterior (purple) sister cells over the final 200 minutes (individual pairs shown as points). Directional bias tested using one-sample Wilcoxon signed-rank tests against zero with Holm correction for multiple comparisons; *** p < 0.001, ns p ≥ 0.05.

We first tested whether the clockwise rotational bias could be explained by the location of new hair-cell pairs within the organ. If such a locational difference existed, it would point towards an effect of the surrounding mechanical environment biasing cell-pair rotations in a specific direction. We examined the dorsoventral distribution of imaged pairs and observed that the CW bias persisted throughout (Figure S10C). Thus, the observed bias does not appear to result from the surrounding mechanical environment.

We next asked whether division angles in *stk32a* mutants were distributed such that they might favor clockwise rotations. However, both WT and mutant neuromasts showed a broad distribution of positive and negative division angles (Figure S10D-E). However, in WT pairs, the rotation direction is correlated with the division angle of the pair (Figure 5D); pairs tended to rotate or align in the direction that minimizes the total rotated angle given their initial orientation. Because only about 50% of WT pairs swap positions across the AP axis, while the remainder tend to execute smaller angular changes, we compared the direction of rotation in these two categories. For a given division angle, the direction of rotation was opposite between aligning and swapping pairs (Figure S10F), reflecting the opposing geometries for cell positioning in these contexts (Figure S10H). On the other hand, in *stk32a* mutants, pairs that divided at a negative angle did not rotate counterclockwise (Figure 5E). Moreover, mutant pairs no longer showed distinct alignment or rotation directions based on the sign of their division angle (Figure S10G).

Therefore, we next asked if *stk32a* perturbation influenced cell movement along the AP axis. However, neither mutation nor transgenic expression of *stk32a* led to pair-wide directional drift over time (Figure 5F). Cell-pair separation, on the other hand, was impacted in *stk32a* mutants (Figure 5G,H). We next looked only at non-swapping pairs, in which it is easier to parse the contributions of individual sisters to their own movements. In WT non-swapping pairs, sisters tend to move in opposing directions, separating over time (Figure 5I, J). While, on average, anterior sister cells in *stk32a*-mutant pairs still migrated anteriorly as in WT, posterior-localized sisters did not migrate posteriorly (Figure 5I,J).

Together, these data suggest that loss of *stk32a* selectively disrupts the directional movement bias of the Notch-ON sister, in which *stk32a* is normally expressed. Notch-ON cells, therefore, do not convert to a Notch-OFF-like behavioral state, but instead may lose their directional bias, mirroring the initially randomized hair-bundle orientation observed in Notch-ON/*stk32a*-mutant cells. Overall, our results demonstrate that Stk32a regulates organ-axis interpretation in the Notch-ON sister and thereby influences both early cell-pair behavior and later hair-bundle polarization, coupling Notch-identity with PCP-provided directional information to drive oriented cell behavior.

We propose a physical, two-dimensional rigid-body model to qualitatively explain the unexpected chiral bias in the mutants (see Methods). By representing the cell pair as two semicircles attached at their shared equator, we can model cytoskeletal movements as forces that generate torque, rotating the cell pair. The direction and placement of these forces on the cell surface are determined by the cell’s Notch state and its *stk32a* expression (Figure 6). In wild-type conditions, we assume that these forces are generated symmetrically on the Notch-OFF and Notch-ON cells (Figure 6A). When the Notch-OFF cell starts in a posterior position relative to its Notch-ON sister, these opposing forces create a robust torque that consistently drives the cell pair to swap positions. Conversely, if the Notch-OFF cell begins anteriorly, the forces pull the cells away from each other into a stable, aligned configuration without inducing a swap. Because the wild-type forces are symmetric, the rotational “tipping point” for the cell pair exactly aligns with the anteroposterior axis, at 0°(Figure 6C). Therefore, the direction in which the pair rotates (clockwise versus counterclockwise) depends cleanly on whether its initial division angle is positive or negative relative to the AP axis. However, the pronounced clockwise chirality observed in *stk32a* mutants suggests a break in the symmetry of these driving forces. Our model proposes that, as a result of *stk32a* loss in the mutant, the force in the Notch-ON cell loses its ventral bias (Figure 6B). Because torque generation is no longer symmetric, the rotational tipping point shifts away from 0°. As a result, the threshold angle for cells rotating clockwise or counterclockwise is no longer aligned with the AP axis, but instead at (*α*_2_ − *α*_1_)*/*2. The shifted tipping point expands the range of initial division angles that result in a clockwise rotation. This result recapitulates our observations: *stk32a* loss-of-function specifically disrupts the balanced force distribution in the Notch-ON sister, biasing the cell pair to preferentially rotate clockwise from a wider range of initial starting angles (Figure 5C,E). It should be noted that the data in Figure 5E suggest that *stk32a-*mutant pairs dividing at negative angles may almost entirely fail to rotate counterclockwise. This behavior is not currently accounted for in our model, and would suggest additional assumptions, for example, that Notch-ON cells become unable to produce force, or that forces in these cells are produced in random directions. While we explored these additional scenarios in our simulations, the available live-imaging data are currently insufficient to distinguish between them. Regardless of this shortcoming, our model, with minimal assumptions, can explain the chiral rotation bias and provides a simple mechanical framework to understand how the interaction between signaling and mechanics produces a robust positional segregation of Notch-ON and Notch-OFF cells. Furthermore, our model hints at a potential role for *stk32a* in determining the subcellular localization of the force-generating machinery driving hair-cell rearrangement.

**Fig. 6.**
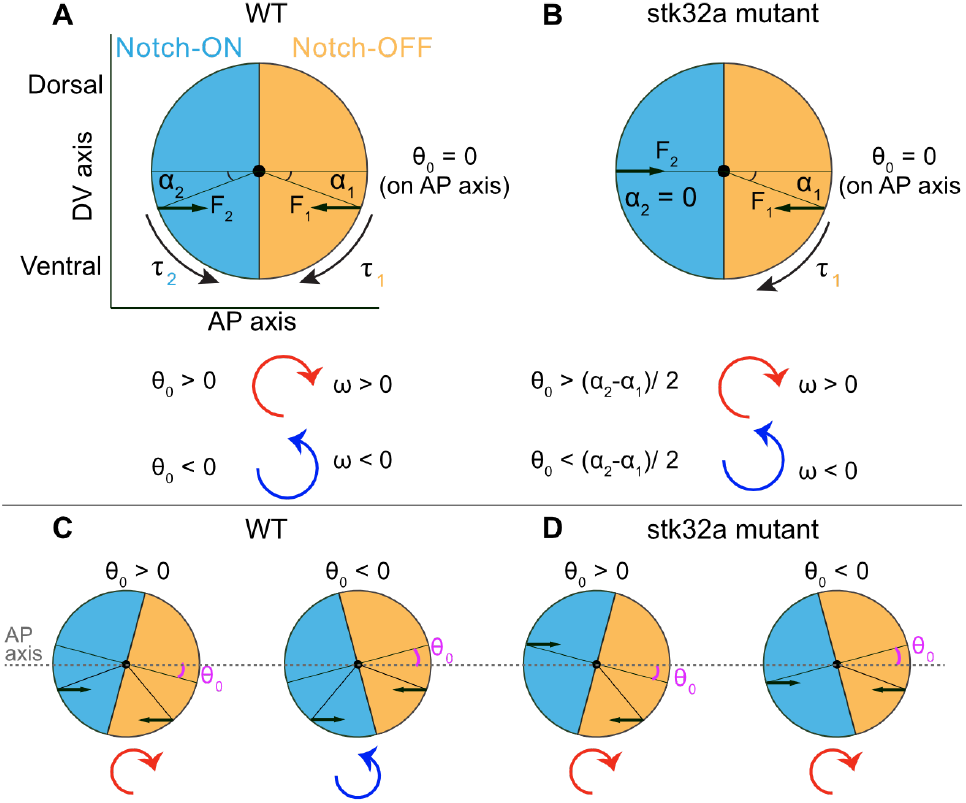
A physical model of cell-pair rotation with a ventral force bias recapitulates the chiral rotation of *stk32a* mutants. **(A)** A WT cell pair in which sister cells are misplaced along the AP axis is shown. The Notch-OFF cell pushes with *F*_1_ and the Notch-ON cell with *F*_2_, and these forces are offset from the midline of the cell by *α*_1_ and *α*_2_ . In this case, rotations are equally likely to proceed in the clockwise or counterclockwise directions (see Methods). **(B)** In an *stk32a* mutant cell pair we assume that *F*_2_ no longer has a ventral bias. Pairs with initial angles *θ*_0_ *>* (*α*_2_ − *α*_1_)*/*2 experience a positive torque and rotate in the clockwise direction. The range of division angles at which cell pairs will rotate CW or CCW depends on the degree of ventral offset in the forces applied by the Notch-OFF sister. **(C)** In WT pairs positive division angles lead to CW rotations (left) while negative division angles lead to CCW rotations (right). **(D)** The predicted loss of ventral bias in the Notch-ON cell of *stk32a* mutants leads to an expanded range of division angles that result in CW rotations.

## Discussion

During development, molecular fate decisions must be translated into physical organization: cells often need to move, remodel, or orient themselves as they differentiate, and must do so in a complex environment. These processes are paradigmatically on display during neuromast morphogenesis. In neuromast hair-cell pairs, Notch signaling establishes opposite polarity identities between sisters that ultimately orient individual hair bundles in mirror-image directions. Our findings support a three-step model: Notch1a first assigns identity, the kinase Stk32a then plays a key role in interpreting that identity to drive directed cell movement, and only after these positional rearrangements do intracellular mechanisms orient the hair bundle. Critically, without *stk32a*, identity is assigned normally but cannot be reliably translated into behavior – the Notch-ON sister fails to execute the polarity-specific movements and generate the bundle orientation that mirror symmetry requires.

Our study resolves conflicting reports on whether Notch1a asymmetry actively drives hair cell-pair rotation (10, 13, 14, 20, 21) or merely buffers stochastic variation (21). By reanalyzing mutants carrying the *hzm17* allele used in prior work (21), we find that rotation frequency is reduced and that pairs exhibit the anteriorly-directed drift previously reported for an independent allele (10), indicating that differences in how swap events were scored, rather than allelic differences, account for prior discrepancies. This highlights the importance of tracking these complex cell movements in three dimensions, which our semiautomated pipeline enables. Our data are consistent with a model in which robust rotations arise because Notch-OFF and Notch-ON cells move in opposing directions at comparable speeds. In this conceptual model, young cells are initially symmetric, and Notch-mediated lateral inhibition breaks this symmetry stochastically. Then, the coupling of cell movements to their Notch status ensures that cell pairs localize in the same configuration, regardless of their initial conditions. This symmetry-breaking model stands in contrast with alternative models in which rotations are driven by one cell type dominating (20). Indeed, our data shows that hair-cell-specific NICD expression drives posterior movement in the opposite direction to *notch1a*-mutant cells, confirming that Notch state cell-autonomously specifies movement direction (Figure 2C). NICD expression disrupts movements more severely than *notch1a* loss, but this likely reflects the earlier onset and higher signaling levels induced by transgenic expression, consistent with elevated *stk32a* transcript levels in *Tg(myo6b:NICD)* relative to endogenous Notch-ON cells (Figure S8D). Rare swaps that occur in Notch-ON or Notch-OFF pairs may reflect cases where sisters moving in the same direction apply unequal forces, allowing one to overtake the other.

A central advance of this work is the identification of Stk32a as a molecular interpreter that converts Notch-dependent fate asymmetry into directed mechanical behavior. We show that Stk32a acts downstream of Notch in the Notch-ON sister to drive polarity-specific movement and rostrad hair-bundle polarization, thereby linking two polarization processes – cell-pair rotation and hair-bundle orientation – that occur at distinct developmental stages and were not previously known to share common regulatory machinery (20). Moreover, we show that in *stk32a* mutants, the initial hair-bundle misorientation is partially corrected over time (Figure S7), suggesting that additional mechanisms exist to ensure the correct polarization of the organ and that hair-bundle polarity remains plastic after it is initially set.

To identify the transcriptional basis of these polarized behaviors, we generated a polarity-resolved transcriptome of hair cells across maturation, separating Notch-ON and Notch-OFF trajectories. Beyond identifying *stk32a* as a Notch-ON-enriched kinase, this dataset reveals genes that define each polarity class and has the potential to elucidate how polarity-specific programs change as cells mature and to uncover the machinery underlying polarity-specific innervation (34–37) and activity (38, 39). Because Emx2 also regulates polarity reversal in the mammalian inner ear (16, 17, 40, 41), this resource further offers a foundation for evolutionary comparisons between the lateral line and the mammalian vestibular system and will enable construction of a more comprehensive gene regulatory network for hair-cell polarity specification more broadly.

Although cell-pair rotations do not occur in the mouse vestibular system, hair bundle polarization is highly conserved (13–16). Our findings confirm that *stk32a* perturbation disrupts zebrafish hair bundle polarization in a manner similar to mouse mutants (31), supporting a conserved role for Stk32a in interpreting planar cell polarity (PCP) cues downstream of fate specification. In mice, *Stk32a* expression in Emx2-negative cells prevents apical localization of the G protein-coupled receptor GPR156 (15, 31, 42, 43), which is expressed in all hair cells but required for polarity reversal in Emx2-positive hair cells (15). In mice, loss of *Stk32a* causes ectopic GPR156 localization at the apical surface of Emx2-negative hair cells, where it fails to planar-polarize, and hair bundles show randomized polarization (31), defects that can be rescued by removal of GPR156 (44). Although GPR156 has yet to be visualized in zebrafish neuromasts due to lack of suitable antibodies, *gpr156* loss-of-function similarly disrupts polarity reversal in fish (15). Consistent with these observations, *stk32a* perturbation in zebrafish randomizes hair-bundle orientation in the Notch-ON/Emx2-negative population (Figure 4L), paralleling findings in the mouse. Together, these results strongly suggest a conserved mechanism in which Stk32a is required for proper interpretation of tissue-wide polarity cues, with additional polarity-specific factors acting in parallel, as proposed in the mouse (31, 44). Future work will be required to confirm this mechanism and to identify substrates (45) and polarity-specific partners of Stk32a. Our polarity-resolved transcriptomic dataset will constitute an important asset in these future endeavors.

*stk32a* mutants show two predominant alterations to their rotation dynamics. First, the percentage of swaps is reduced, but not to the point observed in the *notch1a* mutants or NICD-expressing pairs. Second, the decrease in rotations appears to be driven by an inability of cell pairs to swap in a counterclockwise direction. The clockwise bias in cell-pair rotations suggests a chiral breaking of symmetry in the forces that drive cell movements. We propose that this is driven by a dorsoventral bias asymmetry in these forces (Figure 6), which means that forces are not generated equally throughout cells along the dorsoventral axis, but that a force gradient could exist along this axis. This asymmetry was revealed only through the *stk32a* mutant condition, suggesting that it may be normally masked in WT pairs due to their balanced, opposing forces.

Taken together, our findings argue that tissue organization is not passively inherited from molecular fate but actively constructed through a cascade of mechanically biased behaviors. Notch signaling establishes polarity identity, Stk32a converts that identity into directed cell movements that organize tissue geometry, and intracellular mechanisms then orient hair bundles within this emergent framework. These results support a broader view in which spatial organization during development arises from the progressive translation of fate into behavior, and in which molecular interpreters such as Stk32a play a central role in converting asymmetric signaling into robust, collective polarity patterns.

Ultimately, the morphogenesis of the zebrafish lateral line serves as a powerful model for understanding how molecular fate decisions are actively translated into physical tissue organization. By outlining a stepwise process through which Notch signaling establishes distinct cellular identities, and Stk32a acts as a molecular interpreter to drive directed movements, this work illuminates a critical link between genetic fate and physical morphogenesis. Understanding these intricate morphogenetic rules provides a foundational framework for exploring how complex mechanochemical systems build and refine tissue structures during organogenesis. As the field continues to bridge the gap between transcriptomic identities and three-dimensional cellular behaviors, the resources and mechanistic principles detailed here offer a valuable stepping stone for broader investigations into the evolution and developmental dynamics of highly organized sensory epithelia.

## Methods

### Zebrafish husbandry

Zebrafish experiments were performed according to the guidelines of the Rockefeller IACUC review board (approved protocol #22081-H). In brief, zebrafish were bred in small groups or pairwise crosses, and embryos were raised at 28.5°C in 1× E3 medium (5 mM NaCl, 0.17 mM KCl, 0.33 mM CaCl_2_, 0.33 mM MgSO_4_, 6.25 mM HEPES), adjusted to a final pH between 7.05 and 7.1 and supplemented with 1 *µ*g/mL methylene blue. Experiments were performed using embryos or larvae 2-5 days post-fertilization (dpf). Sex was not determined, as zebrafish do not exhibit sexual differentiation at the larval stages used in this study. WT TU and TL lines, obtained from ZIRC, were used. The published transgenic lines *Tg(myo6b:actb1-EGFP)* (46), *Tg(cldnb:lyn-mScarlet)* (47) and the published mutant line *notch1a(hzm17)* (14, 21) were used. The transgenic lines *Tg(myo6b:NICD-P2A-mCherry, myl7:mScarlet)*^*ru1013Tg*^, *Tg(myo6b:stk32a-EGFP, cryaa:EGFP)*^*ru1014Tg*^ and the mutant line *stk32a(ru800)* were generated in this study.

### Transgenic and mutant strain generation

To generate *Tg(myo6b:NICD-P2A-mCherry, myl7:mScarlet)*^*ru1013Tg*^, referred to as *Tg(myo6b:NICD)* in the text, a geneBlock (ordered from IDT) encoding the Notch1a Intracellular Domain (NICD) (comprising amino acids 1745–2437; RefSeq: NM_131441.1) with a Kozak sequence and attB sites was cloned into pDONR221 to create pME-NICD. This was assembled with p5E-myo6b (46) (from Katie Kindt), p3E-2A-mCherry (AddGene plasmid 26031; RRID:Addgene_26031) (48), and a modified pDEST-395 destination vector (49), replacing GFP with mScarlet – for red heart expression – by Gateway LR recombination. For Tol2 transgenesis, 48 pg plasmid and 100 pg Tol2 mRNA were injected into one-cell embryos. Several founders with germline expression were obtained.

We note that mCherry was not visible in hair cells and transgene transmission was low (2–10% positive F1 progeny). A line showing consistent rostrad-polarity bias in hair bundles was established (Figure 4F and Figure S7F). Time-lapse experiments were performed primarily with F2 embryos, and hair bundle measurements and HCR experiments (Figure S8) were performed with F3 embryos. *Tg(myo6b:NICD)*; *stk32a*^*ru800/ru800*^ analysis was performed with the F4 generation of this transgene.

To generate *Tg(myo6b:stk32a-EGFP, cryaa:EGFP)*^*ru1014Tg*^, referred to as *Tg(myo6b:stk32a*), we ordered a geneBlock (IDT) synthesis of a Kozak sequence followed by the cDNA sequence of the *stk32a* gene (RefSeq: XM_009295698.3, accessed Feb 2024) and flanking attB sites. This geneBlock was cloned into pDONR221 and the resulting pME vector was combined with p5E-myo6b (46) (from Katie Kindt), p3E-EGFP (50) (Addgene plasmid #140877; RRID:Addgene_140877) and pDESTTol2pACryGFP (51) (Addgene plasmid #64022; RRID:Addgene_64022) to generate the injected destination vector. 30 pg of plasmid, along with 50 pg of Tol2 mRNA were injected and a single founder was isolated out of five surviving F0s screened; a line was established from this founder. Experiments were performed with F2 embryos, except the rescue of the *notch1a*^*hzm17/hzm17*^ mutant, which was performed with F3 embryos.

The *stk32a(ru800)* mutant was generated by CRISPR-Cas9 targeting exon 6 of the *stk32a* locus with the sgRNA 5’-CAGCTATGTTGAAATCTGTCAGG-3’. The chemically modified Alt-R CRISPR-Cas9 sgRNA (IDT) was diluted to 7.125 µM in Duplex Buffer (IDT) and combined at equimolar ratio with Alt-R S.p. HiFi Cas9 Nuclease V3 (7.125 µM, IDT) diluted in Cas9 working buffer (20 mM HEPES pH 7.5, 150 mM KCl in nuclease-free water). The ribonucleoprotein complex was assembled by incubating at 37°C for 5 minutes followed by cooling on ice, and 2 nL was injected into the yolk of one-cell embryos. F0 adults were crossed to wild-type fish, and a founder with a high degree of germline transmission of a 4-bp deletion was identified. The deletion results in a frameshift mutation that directly disrupts the predicted DFG motif (52), a critical domain of the AGC kinase family (53). The founder was outcrossed to TL fish, and F1 or F2 heterozygotes harboring the same allele (*ru800*) were incrossed to generate F2 or F3 embryos for experiments.

### Mutant line genotyping

*notch1a(hzm17)* – Genotyping was first performed by amplifying the region surrounding the indel with forward primer 5’-GCTTCTCCTGTAACTGTCCGGC-3’ and reverse primer 5’-CAATGTCGTTTTCACATAGAGCTCC-3’ followed by Sanger sequencing to identify the zygosity of the indel, or through a validated genotyping assay developed at Transnetyx. Prior to live imaging and scRNA-sequencing experiments, mutant animals were phenotyped by screening for somite defects (21, 54).

*stk32a(ru800)* – Genotyping was performed with PCR amplification of the region surrounding the indel using forward primer 5’-GCATAATAGGACCTCTTTACAACCAG-3’ and reverse primer 5’-CCCTTTAACTGAAGTTGTGCTG-3’ followed by Sanger sequencing, or through a genotyping assay (Transnetyx).

### F0 CRISPR screening

To generate crispants, we adapted a previously described protocol for F0 CRISPR screening in zebrafish (32). Briefly, three to four guide RNAs (gRNAs) were designed per target gene, with each gRNA targeting a different exon, preferentially early exons in multi-exon genes. Guide selection was performed using CHOPCHOP (33), prioritizing those guides with high on-target efficiency and minimal off-target predictions. Chemically modified Alt-R™ CRISPR-Cas9 sgRNAs (IDT) and Alt-R™ S.p. Cas9 Nuclease V3 (IDT) were used to generate ribonucleoprotein (RNP) complexes. Each RNP was assembled by combining 28.5 *µ*M sgRNA (in IDT Duplex Buffer) with 28.5 *µ*M Cas9 protein (in Cas9 working buffer: 20 mM HEPES pH 7.5, 150 mM KCl in nuclease-free dH_2_O), followed by a 5-minute incubation at 37°C and cooling on ice. Three to four RNPs targeting the same gene were pooled at equimolar ratios, maintaining a final RNP concentration of 28.5 *µ*M regardless of the number of guides employed. A 2 nL volume of this mixture was injected into the yolk of one-cell stage embryos, delivering approximately 4700 pg of Cas9 protein and 1000 pg total gRNA per embryo (32).

### Time-lapse imaging for cell tracking

Zebrafish larvae (2–2.5 dpf) were anesthetized for ≥15 min in buffered 600 *µ*M tricaine methanesulfonate (Syndel or Sigma) prepared in 1× E3 medium. Larvae were then mounted in 1 % UltraPure low-melting-point agarose (Invitrogen), in 1× E3 supplemented with 600 *µ*M tricaine, on 35 mm MatTek dishes with No. 1.5 coverslips. Larvae were oriented laterally so that the right side of the body lay flush against the coverslip for imaging. Once the agarose had set, fresh anesthetic solution (1× E3, 600 *µ*M tricaine, 1 mM L-ascorbic acid as an antioxidant) was added. Neuromasts were imaged on either an Olympus IX81 or a IX83 microscope equipped with a microlens-based, super-resolution confocal system (VT iSIM, VisiTech International) using a 60x silicon oil (SiOil) immersion objective. During imaging, a temperature-controlled chamber (OKO Labs) was set to 28°C. Images were acquired using either the ORCA Quest qCMOS camera or the Orca ER CCD camera (Hamamatsu Technologies). One to four posterior lateral line neuromasts of the type L1-L4 (4), which are polarized along the anteroposterior axis, were imaged on each fish with time intervals of 3 minutes. For each time point, 64 axial slices spaced 300 nm apart were acquired to capture a volume of 53.6 *µ*m x 53.6 *µ*m x 19.2 *µ*m. To keep regions of interest in frame, we used a custom tracking algorithm to recenter the image at each frame by moving the stage (adapted from https://github.com/a-jacobo/AutoTracker (10)). Fluorescence in *Tg(cldnb:lyn-mScarlet)* was excited by a 561 nm laser and captured through a 605 nm emission filter. *Tg(myo6b:actb1-EGFP)* was imaged either at the first and last time point of the time-lapse, or once every 3 hours using a 488 nm excitation laser and 525 nm emission filter.

Larvae were imaged for a maximum of 48 hours to determine the final polarity of the hair cells imaged, but only cell pairs that divided by 24 hours and had clearly polarized hair bundles were analyzed. Cell pair movements were only analyzed in (1) *notch1a* mutants if both cells were caudad-polarized, (2) *Tg(myo6b:NICD)* transgenic fish if both cells were rostrad-polarized when matured and (3) WT fish if one sister was clearly caudad-polarized and its sister cell rostrad-polarized. Rare cases where neuromasts exhibited abnormal levels of cell death were excluded.

Datasets were generated for WT larvae, *notch1a(hzm17)* mutant larvae, *Tg(myo6b:NICD)* transgenic larvae, *stk32a(ru800)* mutant larvae and *Tg(myo6b:stk32a)* transgenic larvae (with cell-pair sample size ranging from 43 to 75 cell pairs – see respective figure legends). Datasets were generally generated with two fish for a given condition being imaged at a time, depending on embryo availability. For this reason, and the time-intensive nature of these experiments, we created a single WT dataset, which was used as control against the other conditions, and replotted for comparison in Figures 2 and 5.

### Preprocessing, 3D segmentation and cell tracking

A semi-automated pipeline was generated to track three-dimensional metrics and trajectories of cell pairs over time. First, 3D image stacks of neuromast cell membrane signal (*Tg(cldnb:lyn-mScarlet)*), one for each time-point, were deconvolved and bleach-corrected using Huygens Professional (version 24.04). Each image was typically 700×700×64 pixels, with a lateral resolution of 76.6 nm/pixel and an axial resolution of 300 nm/pixel and approximately 800 such images per time-lapse. A custom three-dimensional CellPose 2.0 model was trained on ten hand-segmented deconvolved images of 2 dpf neuromasts using the CellPose Python API (version 2.2) (55). The training data included both lateral and axial sections of the acquired volumes. The custom-trained model was applied to segment images, one per timepoint, in 3D.

Following segmentation, cell masks were tracked across timepoints using particle tracking methods in the Python package Trackpy (version 0.5.0) (56). In general, a search range of 3 pixels and memory of 10 timepoints was employed, with Trackpy’s adaptive search used when tracks were lost beyond memory limit. After tracking was completed, the image stacks of individual timepoints were concatenated to generate a tracked 4D dataset. For every time-lapse, additional images of hair cells, as marked by *Tg(myo6b:actb1-EGFP)*, were acquired at least at the first and last time point. These additional images were used in conjunction with the time-lapse movies to manually annotate the emergence of new pairs of hair cells. 4D segmented masks were manually inspected in Napari (57); segmentation and tracking errors in nascent hair cell pairs were corrected and datasets with persistent tracking or segmentation faults were excluded from further analysis.

### Analysis of cell centroids and angles in three-dimensions

Following manual identification of nascent hair-cell pair segmentations, we extracted cropped image stacks of each pair together with the three-dimensional centroids of the two cells, scaled to account for anisotropic sampling in the z-dimension. Centroid coordinates were rotated to align the organ axis to the x-axis, which corresponds to the anteroposterior axis, and the y-axis to the dorsoventral axis. Additional correction for angled mounting in 3D was unnecessary, as neuromasts were generally mounted flat, so the z-axis corresponds to the apicobasal axis of the neuromast. Because neuromasts were imaged with the fish mounted laterally, the XY imaging plane corresponds to the anatomical sagittal plane (running anterior-posterior and dorsal-ventral), while the z-axis corresponds to depth along the left-right axis of the fish.

The position of cell 1 was defined as *p*_1_ = (*x*_1_, *y*_1_, *z*_1_) and the position of cell 2 as *p*_2_ = (*x*_2_, *y*_2_, *z*_2_). The vector connecting them is defined as (*v*_*x*_, *v*_*y*_, *v*_*z*_) = *p*_2_ − *p*_1_. To assign a consistent orientation to this vector across time and across cell pairs, cell 1 was defined as the more anteriorly positioned cell. This assignment was made at the division timepoint if the intercellular vector was then aligned within 45°of both the AP axis and sagittal plane, or otherwise at the first subsequent timepoint satisfying this criterion. In rare cases in which these criteria were not met, the position at division was used. Then, to calculate the angle *θ* that the cell pair makes to the AP axis, we used the following formula:

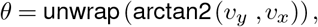

We used the four-quadrant inverse tangent function (*arctan2*) to measure the angle of each cell pair relative to the AP axis at each time point. The resulting angles were then unwrapped to eliminate discontinuities that occur when values cross the 180°boundary. Rotational dynamics were analyzed by comparing the angle at each time point to the angle at the division time point. In image coordinates, where the origin is at the top left corner of the image, the x-axis increases posteriorly and the y-axis increases ventrally, positive values correspond to clockwise rotations and negative values to counterclockwise rotations.

To quantify the elevation angle of the cell pair, *φ*, to the sagittal (XY) plane, we used the following formula:

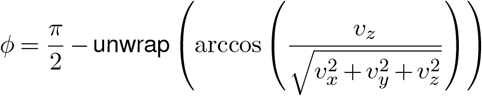

where (*v*_*x*_, *v*_*y*_, *v*_*z*_) represents the vector connecting the two cells. This expression first determines the angle of the cell-pair vector relative to the z-axis (the apicobasal axis), unwraps the values to avoid discontinuities across time, and then converts the measurement to the angle of elevation relative to the sagittal (XY) plane by subtracting from 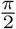.

To identify cell-pair swaps across the anteroposterior (AP) axis, we define two right circular cones with an angle of 45°, and their central axes aligned to the AP axis (Figure S1A). If *θ* and *ϕ* fall within one of these cones, this means that the cell-pair is roughly aligned both to the AP axis and the sagittal plane. A cell-pair swapping event requires two steps: **1**) A cell pair must align to one of the two cones at some time point t_1_, which can be at division or a later time point. **2**) The cell pair must then enter the opposite cone at time t_2_ and remain there for the duration of the time-lapse (up to 10 hours post cell-division).

To calculate deviations of angles from the AP axis (as in Figure 1J), we folded *θ* into the range [0°, 90°], where 0° indicates alignment and 90° indicates maximal misalignment. For deviations from the sagittal plane (as in Figure 1K), we took the absolute value of *ϕ*, which is defined on [-90°, 90°], yielding the same range. We used the angle at division as the initial timepoint and the mean of the last three timepoints of our 10-hour dataset as the final timepoint. To quantify cell-pair reorientation (Figure S1B-C), we plotted the initial division angle against the rotational correction at the final timepoint for each cell pair, where rotational correction is defined as the final angle minus the initial division angle. Both values were folded into [0°, 90°]. Cell pairs were classified as swapping or non-swapping and plotted separately. Linear regression was performed separately for swapping and non-swapping cell pairs to assess whether the degree of rotational correction scales with the initial division angle.

### Static neuromast image acquisition and processing

Static images of neuromasts to assess polarity or expression of genes of interest were also acquired using an iSIM microscope, as described above with the exception that higher laser powers were used to gain a higher signal-to-noise ratio and a 100x SiOil objective was utilized. For visualization in figure panels, brightness and contrast were adjusted on a per-sample and per-channel basis linearly using FIJI. Quantitative measurements (such as HCR *in situ* quantification in Figure S8C-D) were performed on raw, unprocessed data.

### Polarity analysis

The organ’s axis was determined by manually annotating a line connecting the interneuromast cells (as labeled by DAPI and/or *Tg(cldnb:lyn-mScarlet)*) on the proximal and distal sides of the organ of interest. Angle measurements were corrected by rotating measurements and images by this angle. Hair bundle polarity was measured by manually annotating the axis extending from the distal edge of the hair bundle towards the kinocilium and measuring the angle of this line with respect to the organ axis. These measurements were performed at 2 and/or 5 dpf and were restricted to matured hair bundles with clear polarization. Angles were classified into polarity bins spanning the full 0–360° range, with boundaries at 0°, 45°, 135°, 225°, 315°, and 360°. These intervals corresponded to four polarity categories: *Caudad* (0–45°and 315–360°), *Dorsal* (45–135°), *Rostrad* (135–225°), and *Ventral* (225– 315°). Caudad- and rostrad-polarized hair bundles are on-axis and dorsal- and ventral-polarized ones are off-axis. Rose plots were generated using 15°bins with wedge areas scaled proportionally to the fraction of bundles in each bin. In rose plots, bins centered within the off-axis quadrants (45–135°and 225–315°) were highlighted; due to the 15° bin width, highlighted regions span 37.5–142.5° and 217.5–322.5°. Radial axes were scaled independently for each panel.

### Immunohistochemistry

To label Emx2 protein, we adapted a previously reported protocol (14). In brief, dechorionated 2 dpf mutant or transgene-expressing embryos, along with their siblings, were fixed in 4% PFA in 1X PBS at 4°C overnight. Following fixation, embryos were washed four times for 15 minutes each in 0.05% Tween-20 in PBS (PBST) with gentle nutation at room temperature. Embryos were then permeabilized in prechilled 100% acetone at -20°C for 5 minutes and washed 3x 5 minutes in PBST. Blocking was performed for 2-3 hours at room temperature in 1.5% BSA, 1.5% normal goat serum in PBST. Emx2 primary antibody, made in rabbit, was obtained from TransGenic Inc. (product number K0609) and diluted 1:200 in blocking solution. Following blocking, animals were incubated with primary antibody for 24 hours at 4°C. Samples were then washed 4x 15 minutes in PBST and then incubated in 1:250 Alexa Fluor 633 goat anti-rabbit (Invitrogen) in blocking solution overnight at 4°C. Finally, samples were thoroughly washed in PBST, with DAPI (1mg/mL) in the final wash.

To label Vangl2 and spectrin protein, we adapted previously reported protocols (13, 58). In brief, we fixed 4 dpf larvae overnight at 4°C in Prefer Fixative solution with 0.5% Triton added (Anatech). Note that this fixative can quench transgenic fluorophore expression. Following four washes in 1%PBST (Triton) for 15 minutes each, we blocked samples at RT for ∼2 hours in blocking solution (2% Normal Goat Serum, 1% Bovine Serum Albumin, 0.5% Tween-20 in PBS). We then incubated samples overnight at 4°C in 1/100 mouse anti-spectrin *β*II obtained from Santa Cruz Biotechnology (sc-136074), 1/500 rabbit anti-Vangl2 (made against zebrafish C-terminal peptide, gift of Solnica-Krezel laboratory (58)) in blocking buffer. The next day, we washed samples 4x 15 minutes each in 0.1% PBST (Tween-20) and then incubated samples in 1/200 Alexa Fluor 488 goat anti-mouse (Invitrogen), 1/200 Alexa Fluor 568 goat anti-rabbit (Invitrogen) in 0.2% PBST (Tween-20). After overnight incubation at 4°C, we performed four washes in 0.1% PBST, with DAPI in the final wash and left samples in PBS until imaging.

### Hybridization Chain Reaction Fluorescence *in situ* Hybridization (HCR RNA-FISH)

Hybridization chain reaction was performed according to the manufacturer’s protocol (Molecular Instruments) (59, 60) with some modification. Embryos were fixed overnight at 4°C in 4% PFA (in PBS) and then dehydrated in 100% methanol at -20°C at least overnight but up to 3 months. Samples were rehydrated with a series of MeOH/PBSTween (0.1%) washes up to PBST(0.1%). Embryos were then treated with 10*µ*g/mL of proteinase K for 3 minutes followed by two quick washes of PBST. Embryos were then post fixed for 20 minutes at RT in 4% PFA in PBS followed by 5 washes in PBST (5 minutes each). Next, embryos were pre-hybridized with prewarmed probe hybridization buffer for 30 minutes to 1 hour at 37°C. Then, 2-4 pmol of probe solution was added to 500 *µ*L of hybridization buffer and samples were incubated overnight at 37°C. The following day, samples were washed 4x 15 minutes in pre-warmed probe wash buffer (at 37°C), followed by two five-minute washes in 5X SSCT at RT. Next, samples were incubated for 30 minutes in RT amplification buffer. The matching hairpin solutions were prepared by separately heating h1 and h2 hairpins to 95°C for 90 seconds, followed by 30 minutes of cooling in a dark drawer at RT. A combination of h1 and h2 hairpins were added to each sample and samples were incubated overnight at room temperature in the dark. The following day, samples were washed 2×5 min, 2×30 min and 1×5 min in 5X SSCT with the last wash containing DAPI (1mg/mL). Samples were stored in 5X SSCT at 4°C until imaging. The following probes were utilized: *stk32a, dlc* and *hey2*. Hairpin amplifiers were conjugated to Alexa546 or Alexa647.

### Embryo dissociation for scRNA-sequencing experiments

scRNA-seq was performed on hair cells isolated from 4 dpf larvae across three conditions: WT, *notch1a(hzm17)* mutants, and their siblings (a mix of WT *(+/+)* and heterozygotes *(hzm17/+)*). WT larvae were obtained from incrosses of *Tg(myo6b:actb1-EGFP)* fish, whereas mutants and siblings were obtained from incrosses of *notch1a(hzm17/+);Tg(myo6b:actb1-EGFP)* fish. Homozygous mutants were identified at 3 dpf by somite defects, allowing separation from siblings. Each condition was replicated twice, with mutant and sibling samples collected from the same embryo batches on the same day. WT replicates consisted of 450–600 larvae, mutant replicates ∼250 larvae, and sibling replicates 250–400 larvae.

Embryo dissociation and FACS was performed using an adapted version of a previously published protocol (25). In brief, 4 dpf larvae were anesthetized in tricaine, separated into batches of 150 larvae and placed in cell strainers in a 6 well plate. Strainers were then moved to a new well containing 4.5 mL ice cold 0.25% trypsin/1X EDTA (Gibco). All subsequent steps were performed either on ice or at 4°C. Larvae were transferred to a GentleMACS C-Tube (Miltenyi Biotec) and the GentleMACS device was used to gently dissociate the tissue, by mixing 240 rpm for 5 minutes, switching the direction of mixing each minute. We did not aim for complete larval dissociation, as neuromasts are at the surface of the skin. Next, the larval bodies were removed by filtering using a 70 *µ*m filter (Filcons) into a 14 mL round-bottom polypropylene tube (Falcon). The tubes, one for each batch of 150 larvae, were then spun down in a swinging bucket centrifuge for 5 min at 2000 rpm. The supernatant was removed, and the cell pellet was washed twice with ice cold 1x DPBS with 0.04% BSA, each with a 5 min spin down at 2000 rpm with slow deceleration. At each wash, the cell pellet was fully resuspended with gentle pipetting using a 1000 *µ*L pipette tip and ∼20 gentle pipetting steps to fully resuspend. Following the second DPBS wash, the DPBS was removed, and each cell pellet was resuspended with 400 *µ*L DPBS, 10 mM HEPES(pH 7.2), 0.04% BSA, 0.2U/*µ*L Protector RNase inhibitor (Roche). Resuspended cell pellets corresponding to the same sample were combined and filtered through a 70 *µ*m Filcons strainer into a polypropylene tube that was precoated with 1% BSA to prevent sticking.

Following resuspension, cells were stained with 4 *µ*M DRAQ5 (ThermoFisher) prior to sorting. DRAQ5+/EGFP+ cells were sorted by the Rockefeller Flow Cytometry Resource Center on the BD FACSAria using a 100-micron nozzle into cell sorting collection tubes containing BSA, HEPES and Protector RNase inhibitor, resulting in a final concentration of ∼0.04% BSA, 10 mM HEPES and 0.2 U/*µ*L Protector RNase inhibitor in the sorted solution. Sorting was terminated at 40 minutes to ensure that genomic analyses were performed on healthy cells. Due to the rarity and fragility of hair cells following sorting, no counting of cells or resuspension was performed. Rather, based on several trial runs, we expected viability to be ∼75-80% and estimated counts of cells to be about 70% of the sorting count.

### 10× Chromium scRNA sequencing

Following sorting, cells were gently pelleted and promptly loaded on the 10x Chromium Controller at the Rockefeller Genomics Core Facility to generate Gel Bead-in-Emulsions (GEMs). cDNA was isolated, amplified, and used to construct sequencing libraries with unique sample indices, generated using the 10x Genomics Chromium Single Cell 3’ Reagent Kits v3.1 (Next GEM). Libraries were pooled at equimolar ratios and sequenced on an Illumina NovaSeq 6000 using S4 v1.5 reagents and Control Software v1.7.5, following the manufacturer’s protocol (Doc# CG000204, Rev.D). Sequencing was performed with paired-end reads (28 bp Read 1, 8 bp i7 index, 91 bp Read 2).

### scRNA-seq read alignment and ambient RNA correction

Sequencing reads were demultiplexed and mapped using CellRanger 7.1.0 to the zebrafish reference genome (GRCz11.109, Ensembl (61)) pre-filtered for protein-coding genes. To address ambient RNA contamination, we applied the *remove-background* module of CellBender (version 0.2.1) (62) to identify and exclude empty droplets and correct for ambient RNA in droplets containing real cells. Each of the six samples was processed separately, initially using 150 epochs, a false positive rate of 0.01, and a default learning rate of 0.0001. Parameters were adjusted as needed to produce clear sigmoidal elbow plots that distinguished empty droplets from those containing cells. For the remaining analyses, we used the CellBender-corrected counts matrices as our data input.

### Preprocessing and quality control of scRNA-sequencing dataset

A Seurat (v4.3.0) (63) object was generated using Cell-Bender (62) filtered cells and corrected counts from all six samples. We merged the samples without any batch correction and performed quality control on the merged object. Genes expressed in fewer than five cells were excluded. To remove potential empty droplets and doublets, we removed cells if they (1) expressed fewer than 2,500 genes, or (2) more than 8,000 genes, (3) had over 45,000 UMIs, or (4) contained more than 5% mitochondrial RNA content. This filtering step resulted in 4873 WT cells, 2475 mutant cells and 2784 sibling cells. The data were then clustered with a standard Seurat pipeline. FeaturePlots and heat maps examining hair cell marker genes, in addition to potentially contaminating cell types were generated to subset the lateral line-specific hair cell clusters. This filtered population was renormalized and clustered and a second round of filtering was performed to remove non-lateral line hair cells still present. The subsequent analyses used this cleaned lateral line hair cell dataset, which contains 2056 WT hair cells, 1364 mutant hair cells and 1100 sibling hair cells. From this data set, young hair cells were selected by choosing cells that express the delta-ligand *dld*, a young hair cell marker (24), and that do not express mature hair cell marker *tekt3*; this dataset was further filtered to remove cells expressing (1) more than 7000 genes and (2) over 35,000 UMIs, to remove remaining potential doublets.

### Differential gene expression analysis on pseudobulked samples

To compare the transcriptomes of *notch1a* mutant hair cells with those of sibling and wild-type (WT) controls, we used a pseudobulk RNA-seq approach (27, 64). Raw counts were aggregated per sample using Seurat’s *AggregateExpression* function, yielding two replicates per condition (mutant, sibling, WT). Sibling and WT replicates were combined into a single control group, and differential expression was performed using DESeq2 (26) with a design comparing *notch1a* mutant to sibling/WT (65). Genes with fewer than 5 total counts across samples were excluded. P-values were adjusted using the Benjamini-Hochberg method. Results were filtered for adjusted p-value < 0.05, base mean expression > 40, and absolute log2 fold change >= 1.

### UMAP embedding to separate hair cell polarities

To identify transcriptional programs associated with polarity specificity, we leveraged the known polarity-specific expression of *emx2* (13, 14, 16). We first performed PCA on the single-cell dataset and calculated the absolute Pearson correlation between each cell’s score across the top 50 PCs and its SCT-normalized *emx2* expression. PCs whose scores covary with *emx2* may reflect transcriptional programs that distinguish *emx2*-positive and *emx2*-negative cells (*i*.*e*., polarity-specific programs). We ranked PCs by their correlation strength versus *emx2* and selected those above the elbow of the ranked correlation profile (Figure S6C,D), retaining the top 4 PCs for the full dataset and top 6 PCs for the young hair cell dataset. From these selected PCs, we identified genes with the highest loading strengths (up to 1000 per PC with minimum absolute loading of 0.05) and selected the union of genes across PCs; for the young hair cell dataset, this procedure yielded 232 unique genes. We then used this gene set as input features for UMAP embedding, using the *runUMAP* function for 5000 epochs to generate a polarity-specific low-dimensional representation of the data.

### Pseudotime analysis and differential expression with Slingshot and tradeSeq

We applied *Slingshot* (66) to infer developmental trajectories and assign pseudotime values within the young hair-cell embedding. Cells were clustered in the polarity-specific UMAP space at a resolution of 0.5, and a root cluster was specified as the starting point for trajectory inference. This analysis identified two distinct lineages and assigned pseudotime values to cells along each lineage originating from the root cluster. We then used the package tradeSeq (67) to identify genes whose expression changed along pseudotime in the two lineages (Notch-OFF and Notch-ON). For this analysis, we ran the *fitGAM* function with the raw counts for the 3000 most highly variable genes (filtered to require at least 10 cells with more than 3 counts) and 4 knots. We then ran the function *patternTest* to identify genes with significantly different expression patterns between lineages (Supplemental Table 3). To generate the heatmap from the *patternTest* output, we kept the top 30 genes that met the criteria (fcMedian > 0.5, FDR-adjusted p < 0.05). Smoothed expression curves were predicted along pseudotime with the TradeSeq function *predictSmooth*, and for heat-map visualization, each gene’s expression profile was independently scaled using min-max normalization to examine relative temporal dynamics across pseudotime.

### Statistical analysis

Data visualization and statistical analysis were performed with Python (v.3.9) and R (v.4.4). The details of statistical testing are given in the appropriate figure legends; p-values were generally corrected using the Holm method.

### Rigid-Body Simulation of Cell-Pair Rotation

To model the mechanical dynamics of cell-pair rotation, we developed a two-dimensional rigid-body simulation. We represent a developing cell pair as a single two-dimensional rigid body composed of two solid semicircles of radius *R*, joined along their shared diameter. Within the body reference frame, the two cells are defined by their distinct cellular identities: Notch-ON and Notch-OFF (Figure 6A). The spatial configuration of this cell pair in the laboratory frame is uniquely defined by its center of mass position, denoted by **q** = (*x, y*) ∈ ℝ^2^, and an orientation angle *θ*. Assuming that both cells have the same mass *m*, the total mass of the system is *M* = 2*m*, and the moment of inertia about this center equals that of a full solid disk, formulated as 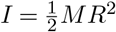.

We model the mechanical forces generated by cytoskeletal movements as fixed point-forces in the laboratory frame, applied directly to the surface of each cell. Specifically, the Notch-OFF cell exerts a force **F**_1_ = (− *F*, 0) at a specific arc angle *α*_1_ relative to its pole, whereas the Notch-ON cell exerts an equal and opposite force **F**_2_ = (+*F*, 0) at an arc angle *α*_2_. Because these applied forces are equal in magnitude and opposite in direction along the x-axis, their translational contributions cancel. As a result, the center of mass of wild-type cell pairs does not translate. In the case of *notch1a* mutant and overexpression experiments, these two forces will point in the same direction, producing a movement of the center of mass (Figure 2C-G).

The contribution of each cell to the total torque can be shown to be

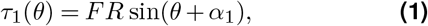

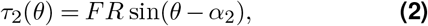

so the total torque is

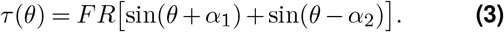

Starting from Eq. (3) we apply the sum-to-product identity 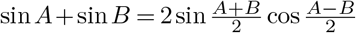 with *A* = *θ* + *α*_1_ and *B* = *θ* − *α*_2_:

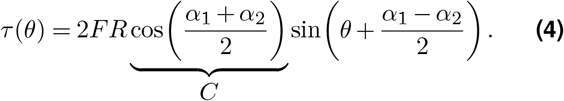

When cells are in the swapping configuration arc angles satisfy |*α*_1,2_| *<* 90 and *C >* 0. Defining the *zero-torque angle*

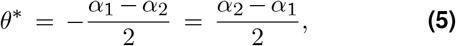

Eq. (4) simplifies to

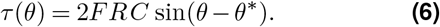

Since 2*FRC >* 0, the sign of *τ* is determined entirely by the factor sin(*θ* − *θ*^∗^). This geometric relationship demonstrates that a symmetric force placement (*α*_1_ = *α*_2_) produces a zero-torque angle of *θ*^∗^ = 0°, resulting in no inherent rotational bias. In contrast, if the force application point on the Notch-ON cell is shifted dorsally relative to the Notch-OFF cell (i.e., *α*_1_ *> α*_2_), it yields a zero-torque angle (*θ*^∗^ *<* 0°). For *θ*_0_ *> θ*^∗^ the resulting torque is positive, producing a CW rotation. This shift creates an unstable equilibrium boundary that inherently biases the cell pair toward clockwise rotations, successfully recapitulating the chiral bias observed in our experimental data.

## Data and Code Availability

Single-cell RNA-seq data have been deposited in GEO (under accession number GSE326142). The original analysis code is on Github (https://github.com/egatlas/cell-pair-rotations-2026), and the processed segmented cell pair data and cell centroid tracking data will be deposited at Zenodo. Due to size restrictions, raw imaging data is available on request.

## Declaration of generative AI in scientific writing

During the preparation of this manuscript, the author(s) used ChatGPT 5.2 and Claude Opus and Claude Sonnet 4, and Gemini 3.1 to edit sections of the text to improve clarity and readability. After using these tools, the author(s) reviewed all edits and take full responsibility for the content of the manuscript.

## Acknowledgments

We would like to dedicate this paper to the memory of A. James (Jim) Hudspeth, thesis advisor of the first author. We thank Samantha Campbell for expert fish husbandry. We are grateful to Anna Erzberger, Francesco Gianoli, Agnik Dasgupta and Xinyue Deng for insightful discussions. We thank Cornelia Bargmann and all members of the Hudspeth Lab for their comments on the manuscript. We thank Rohan Roy, Swarna Jeewajee, Agnik Dasgupta and Gaurav Shrestha for help screening *notch1a* mutants for scRNA-sequencing experiments. We also thank the Rockefeller Flow Cytometry core facility, Genomics core facility and Bioinformatics core facility for crucial expertise in cell sorting, library prep, sequencing and for scRNA-seq analysis training. We thank Daniela Munch and Tatjana Piotrowski for advice on the sample preparation for scRNA-sequencing, Lilianna Solnica-Krezel for the Vangl2 antibody, Katie Kindt for the *myo6b* promoter and Hernan Lopez-Schier for the *notch1a* mutant fish. EA thanks the Fisher Center for Alzheimer’s Research Foundation for support of her graduate work. AJ is grateful to Biohub donors P. Chan and M. Zuckerberg for their generous support. We thank the Howard Hughes Medical Institute for supporting this work.

## Author Contributions

EA, AJ and AJH conceptualized the work. EA designed methodology in consultation with AJ, AJH. EA performed all experiments with the help of SK in optimizing and performing the F0 CRISPR screen. AJ developed the initial version of the 3D tracking and segmentation pipeline; EA further improved it and fine-tuned the segmentation model. EA led data analysis efforts with input from BF, CR, AJ, AJH. CR designed and implemented polarity-specific clustering of the scRNA-sequencing data and BF aided in 3D interpretation of angular trajectories and implemented the cell-pair swapping definition. AJ formulated the mathematical model. EA wrote the initial manuscript and EA and AJ revised the manuscript with input from all authors other than AJH. AJH, AJ, and EA supervised and AJH acquired funding for this project.

## Supplemental Information

**Supplemental Figure S1.**
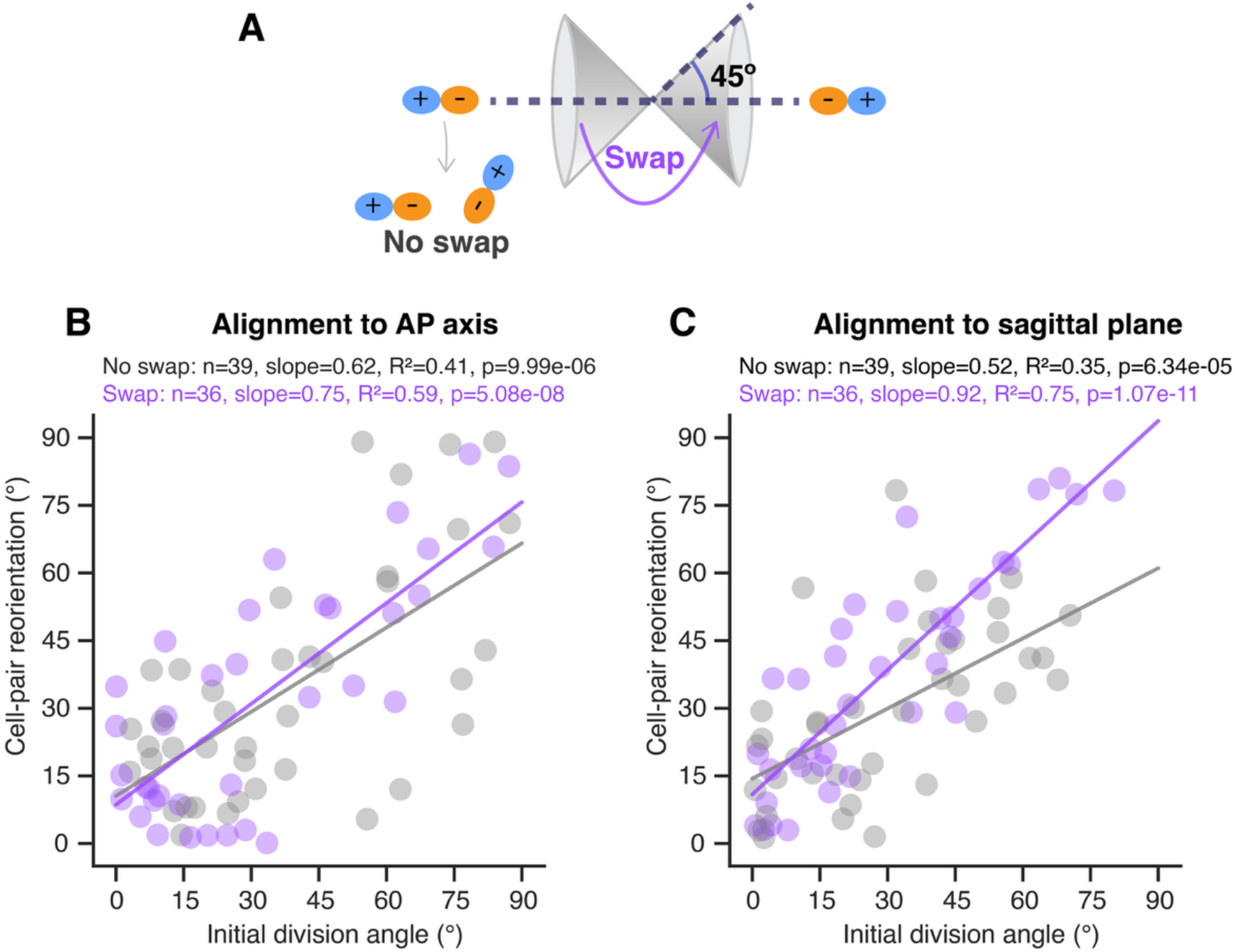
Cell pair behaviors align cells in 3D, related to Figure 1. **(A)** Schematic depiction of cell-pair swapping definition (see Methods). Cell pairs that align within 45º of a cone around the AP axis and end in the opposite cone are defined as swapping pairs. **(B-C)** Scatter plots showing the division angle with respect to either the AP axis **(B)** or angle of elevation from the sagittal plane **(C)** as compared to the degree of cell-pair reorientation, folded to a range from 0-90º. There is a positive correlation between initial division angle and the amount of correction. Each point represents a single cell pair, colored by swapping status (swapping: purple, non-swapping: gray). Linear regression analysis was performed separately for swapping and non-swapping pairs, and separate regression lines and accompanying statistics are shown on each plot.

**Supplemental Figure S2.**
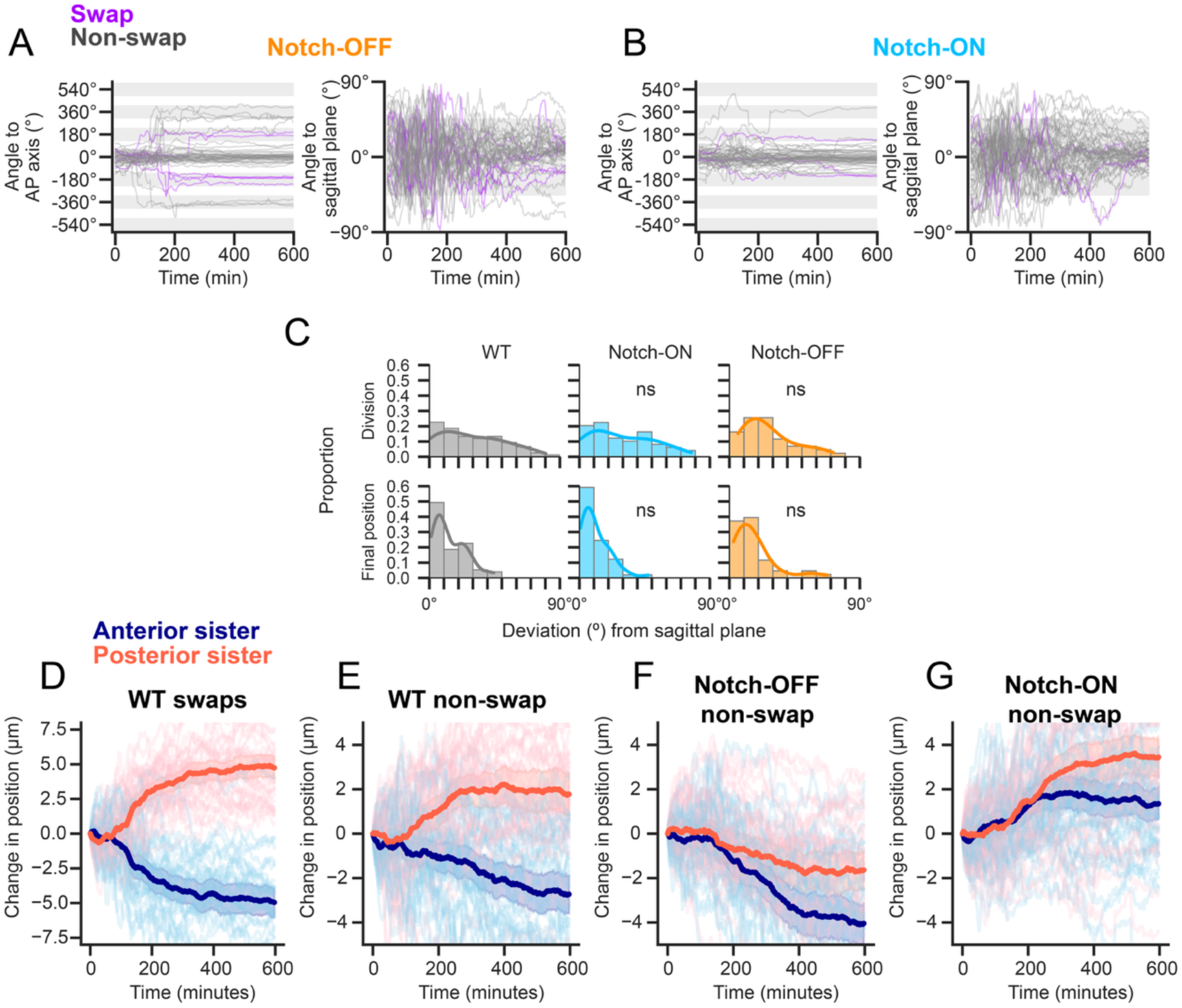
Extended data from Figure 2. **(A-B)** Cell-pair angular trajectories with respect to AP axis and sagittal plane for **(A)** Notch-OFF and **(B)** Notch-ON pairs. Gray bands represent ±45º around the AP axis. The trajectories of swapping pairs are shown in purple and non-swapping pairs in gray. **(C)** Deviation of all cell pair angles from the sagittal plane at timepoints of division and at the end of the timelapse. KS test with Holm correction tested whether spreads of Notch-OFF or Notch-ON pair deviations differed from WT at each timepoint; ns, p ≥ 0.05. **(D-G)** Plots of individual cell centroid movements over time. For each pair, the cells that ended posteriorly (in red) and anteriorly (in blue) were grouped, and their average position was calculated over time. Error bands represent the 95% CI. **(D)** WT pairs that undergo swaps (n=36 pairs). Note that the axis limits here are larger than in subsequent panels, as swapping pairs undergo greater positional shifts. **(E)** WT non-swapping pairs (n=39). **(F)** Notch-OFF non-swapping pairs (n=37). **(G)** Notch-ON non-swapping pairs (n=46). Axis limits are truncated to prioritize visualization of the average trajectories, which results in some individual traces extending beyond the displayed range.

**Supplemental Figure S3.**
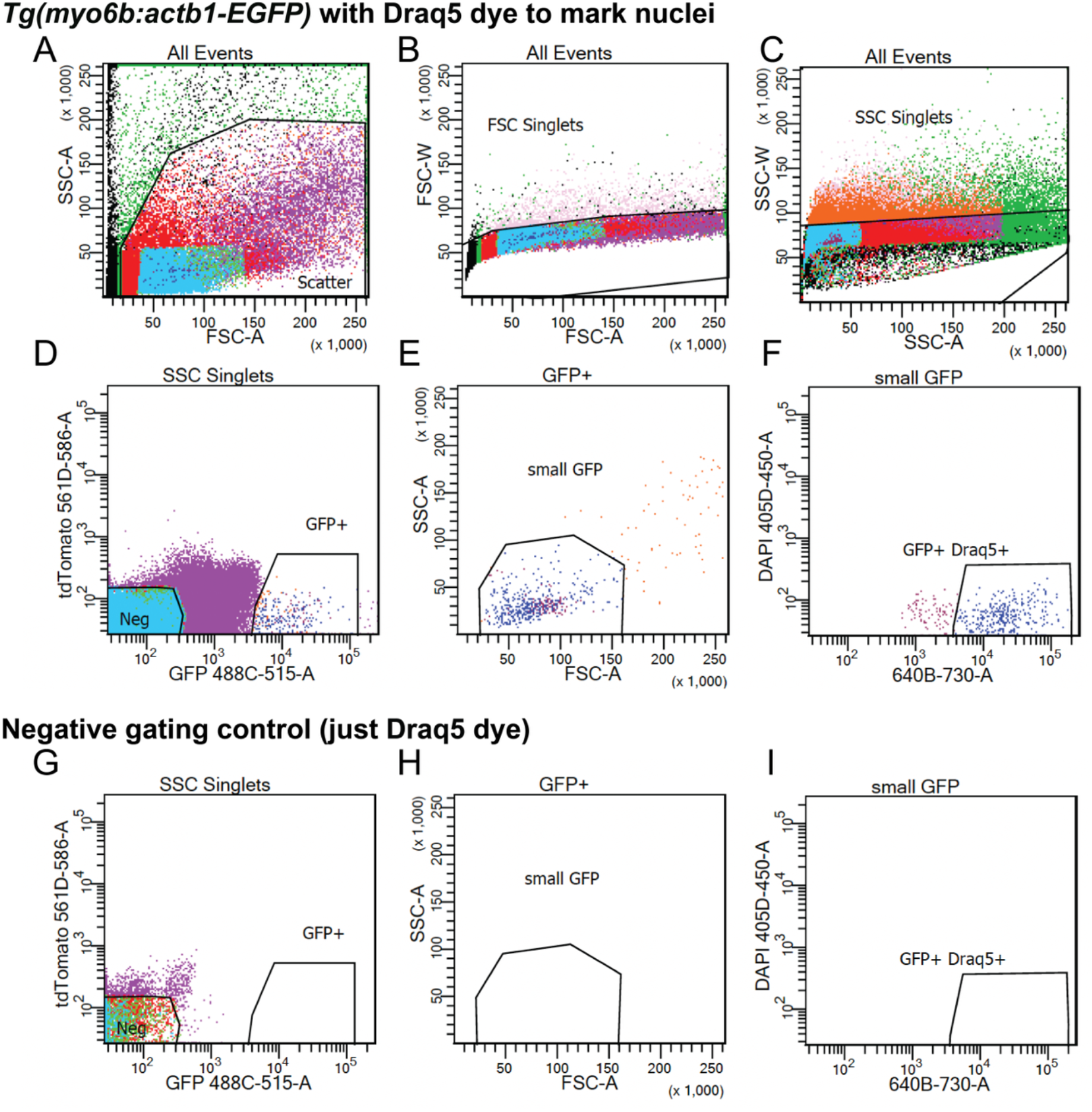
Flow cytometry gating scheme for isolating GFP+ hair cells, related to Figure 3. **(A)** Initial gating on FSC-A vs SSC-A was used to exclude debris, followed by singlet selection based on **(B)** FSC-A vs FSC-W and **(C)** SSC-A vs SSC-W. Following singlet gating, **(D)** GFP vs. tdTomato plots identified GFP+ events (note: cells did not express tdTomato). **(E)** Cells were then gated to select small cells based on FSC-A vs SSC-A, as pilot experiments demonstrated that hair cells are small. **(F)** Draq5 staining was used to select nucleated GFP+ cells, thus excluding cellular debris. A negative control sample stained with Draq5 only (no GFP transgene) was analyzed in parallel **(G-I)** to define the GFP+ gate and exclude autofluorescence. Note that a live/dead exclusion step was not possible because mechanosensory channels in mature hair cells permit the uptake of dyes such as DAPI, leading to the erroneous classification of viable cells as dead^1^.

**Supplemental Figure S4.**
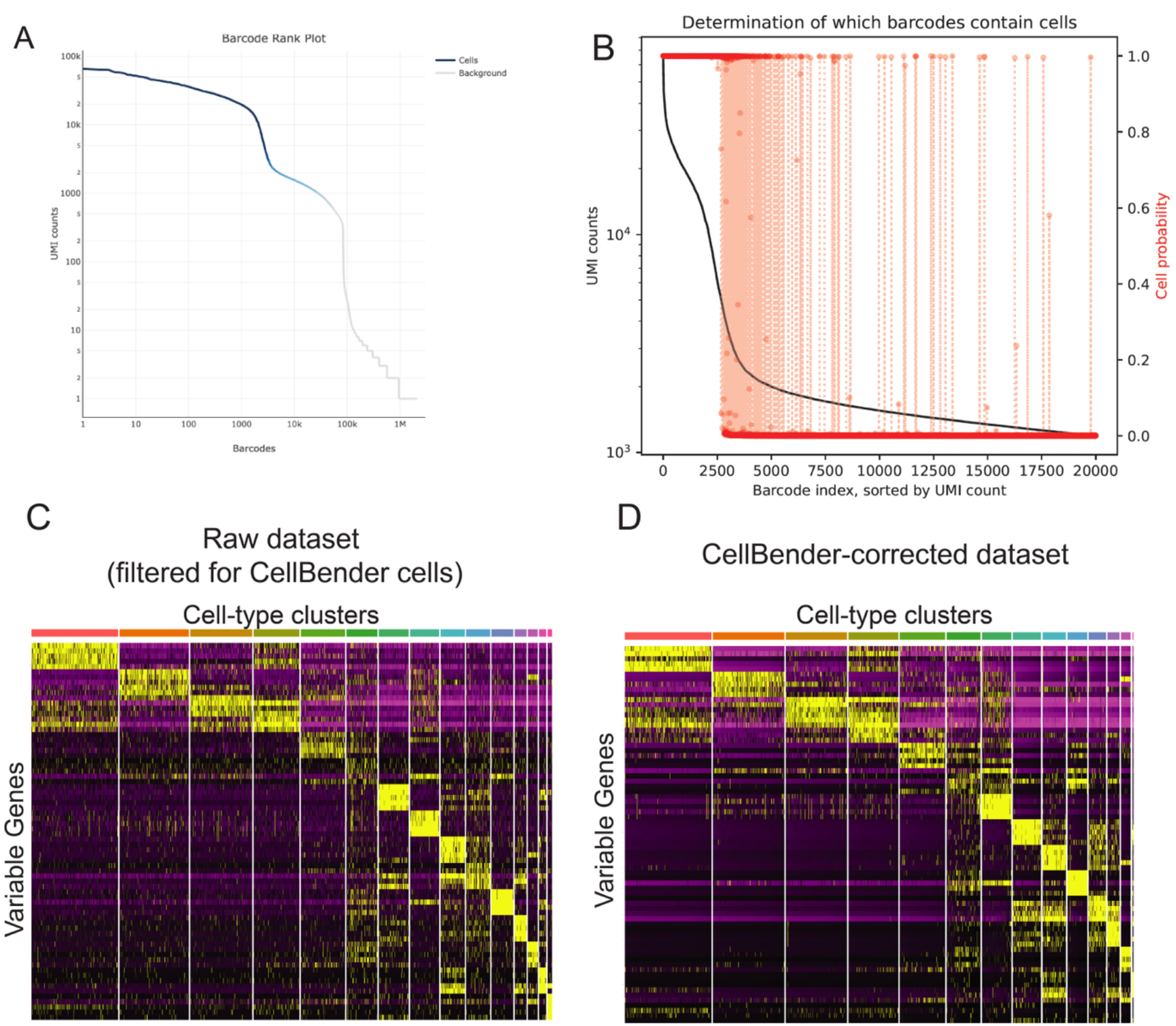
CellBender-aided selection of cell-containing droplets and ambient RNA correction, related to Figure 3. **(A)** CellRanger-generated elbow plot of Unique Molecular Identifier (UMI) counts per droplet barcode (one WT replicate shown as an example). Barcodes are ordered along the x-axis by decreasing total UMI counts per droplet on a log-scale (y-axis). The blue portion of the curve denotes droplets classified as cells, while the gray portion represents droplets identified as empty. **(B)** CellBender-generated elbow plot showing total UMI counts per barcode (black curve, log-scale, left y-axis) versus barcode index sorted by decreasing UMI count (x-axis), overlaid with CellBender’s estimated probability that the droplet contains a cell (red dots, right y-axis). High-UMI barcodes are determined to have a probability near 1 of containing cells, while low-UMI barcodes to the right fall to near-zero probability (empty droplets), demonstrating that the CellBender tool is functioning as expected. **(C)** Heatmap showing gene-wise scaled expression (z-scores), with purple indicating low expression and yellow indicating high expression, for selected variably expressed genes (rows) across single cells (columns). Cells are grouped by cluster identity (top color bar). In the raw CellBender-filtered dataset, many cluster-specific genes show broad expression. **(D)** Same as in panel C, but for CellBender-corrected dataset. Heatmap shows sharper, cluster‐specific expression patterns and cleaner separation between clusters.

**Supplemental Figure S5.**
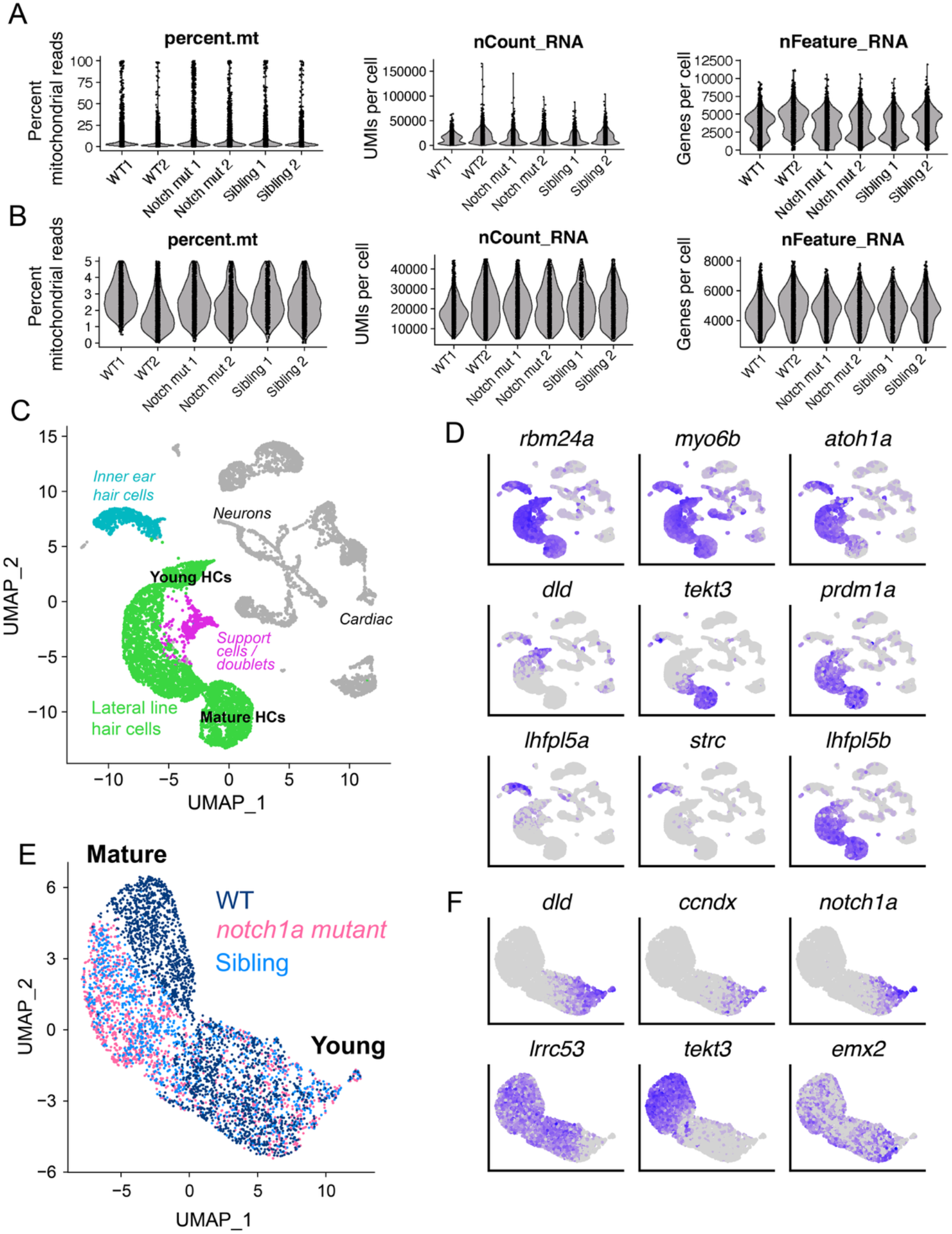
Quality control and initial clustering of lateral line hair cells, related to Figure 3. **(A)** CellBender-corrected count matrices from all six samples (two replicates per condition) were merged prior to quality control. <u>Left</u>: a small population of likely dying cells exhibited extremely high *percent*.*mt* (percentage of mitochondrial reads). <u>Middle</u>: distributions of *nCount_RNA* (number of UMIs per cell) were multimodal. <u>Right</u>: distributions of *nFeature_RNA* (number of detected genes per cell) were multimodal. **(B)** Same as panel A but following the application of quality control thresholds to the dataset. **(C)** A UMAP of all cells in the dataset, with the lateral line hair cells labeled in green. Other contaminating cell types (such as inner ear hair cells, neurons, and cardiac cells) are also shown. A cluster (marked in pink) that had higher counts and expressed both support cell and hair cell markers was also observed and may represent support cell-hair cell doublets. These cells were excluded from further analysis. **(D)** Marker gene expression used to assess the identity of lateral-line hair cells. <u>Top row</u>: hair cell-specific transcripts, expected to be expressed in both lateral-line and inner-ear hair cells. <u>Middle row:</u> *dld* and *tekt3* mark young and mature hair cells^2^, respectively, while *prdm1a* encodes a transcription factor crucial in determining lateral line hair cell fate^3^. <u>Bottom row</u>: Inner ear-specific genes (*lhfpl5a, strc*) and lateral-line specific paralog *lhfpl5b* were used to further distinguish inner ear from lateral-line hair cells^4,5^. **(E)** UMAP dimensionality reduction on filtered set of lateral line hair cells. *notch1a*-mutant cells are marked in pink, their siblings (heterozygous and WT hair cells) in light blue, and the separate population of WT hair cells in navy blue. Young and mature hair cells were identified by the expression of genes shown in F. **(F)** The expression of young and mature hair cell transcripts. Top row: *dld, ccndx*, and *notch1a* mark young hair cells^2,6^. Bottom: row: *lrrc53* and *tekt3* mark more mature hair cells^2^. *emx2* RNA is known to be specific to Notch-OFF hair cells^7,8^ and its expression exhibits a more heterogenous pattern across both young and mature hair cells.

**Supplemental Figure S6.**
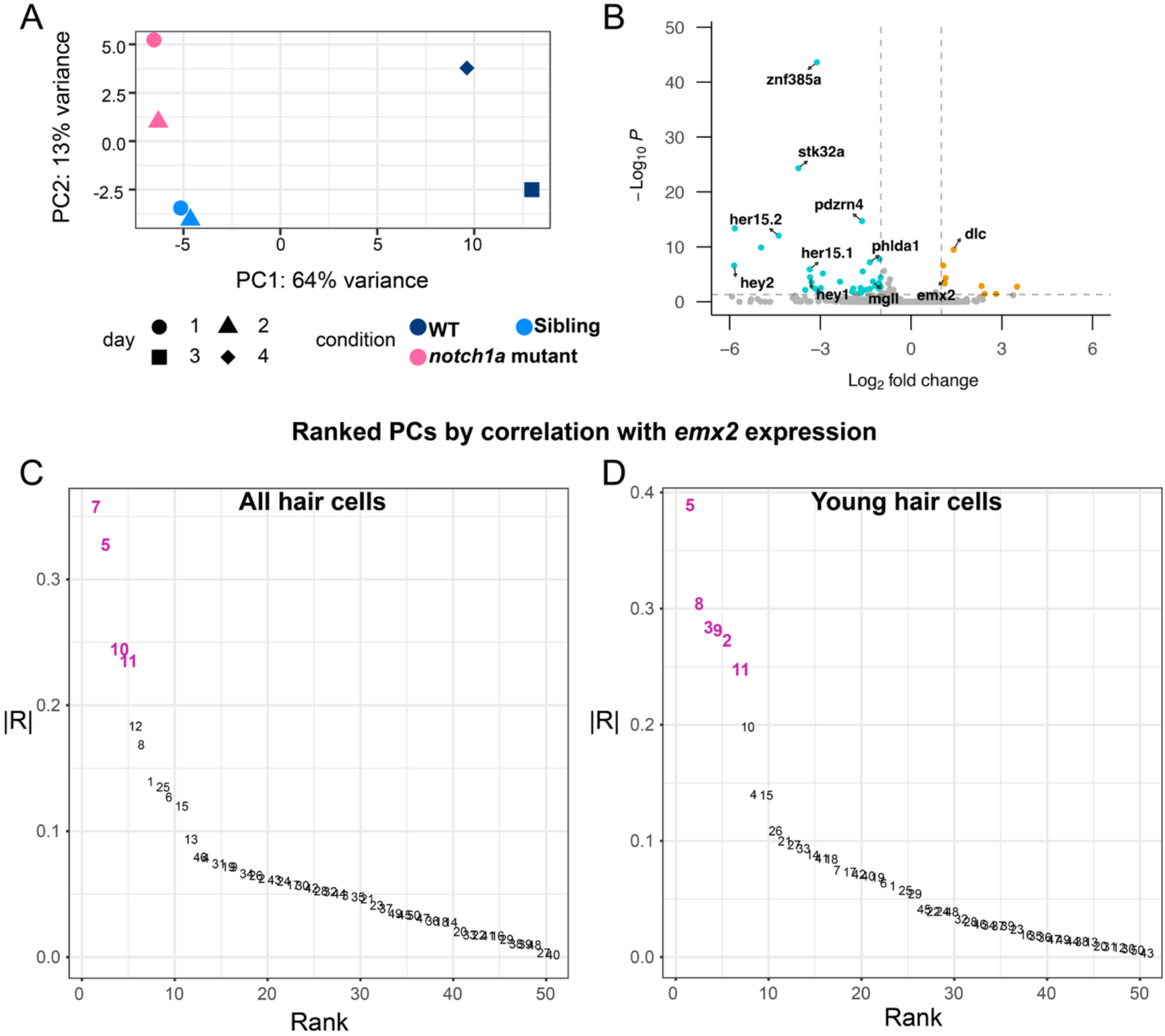
Differential gene expression and search for polarity-specific signature, related to Figure 3. **(A)** Principal component analysis (PC1 vs. PC2) of preprocessed hair cells, separated by sample. Each point is one of the six samples; color denotes genotype, whereas point shape denotes day of collection. PC1 represents 64% of the variance of the data, while PC2 represents 13%. **(B)** A volcano plot showing genes differentially expressed in young (*dld+, tekt3-)* hair cells from *notch1a*^*hzm17/hzm17*^ mutants compared to both siblings, and other wild-type hair cells. The x-axis represents the log_2_ fold change, and the y-axis shows the –log_10_ of the FDR-adjusted *p*-value. Differential expression was assessed using DESeq2. Genes with abs(Log_2_ fold change) > 1 and adjusted p-value < 0.05 are highlighted — blue indicates putative markers of the Notch-ON state, and orange indicates putative Notch-OFF markers. Selected significantly differentially expressed genes are labeled, including known polarity-specific transcription factor *emx2*, and canonical Notch-signaling targets *hey1, hey2, her15*.*1*, and *her15*.*2*, along with Delta ligand *dlc*. **(C)** Elbow plot exhibiting the absolute correlation of *emx2* expression with principal components 1-50 for all hair cells. **(D)** Same analysis as in C for young hair cells only. Each point shows the absolute correlation (|R|) of the given PC scores with *emx2* expression, ranked from highest to lowest. PCs shown in pink had the strongest associations, and the top loadings (*i*.*e*., genes) of these PCs were used to generate a new, polarity-specific UMAP embedding.

**Supplemental Figure S7.**
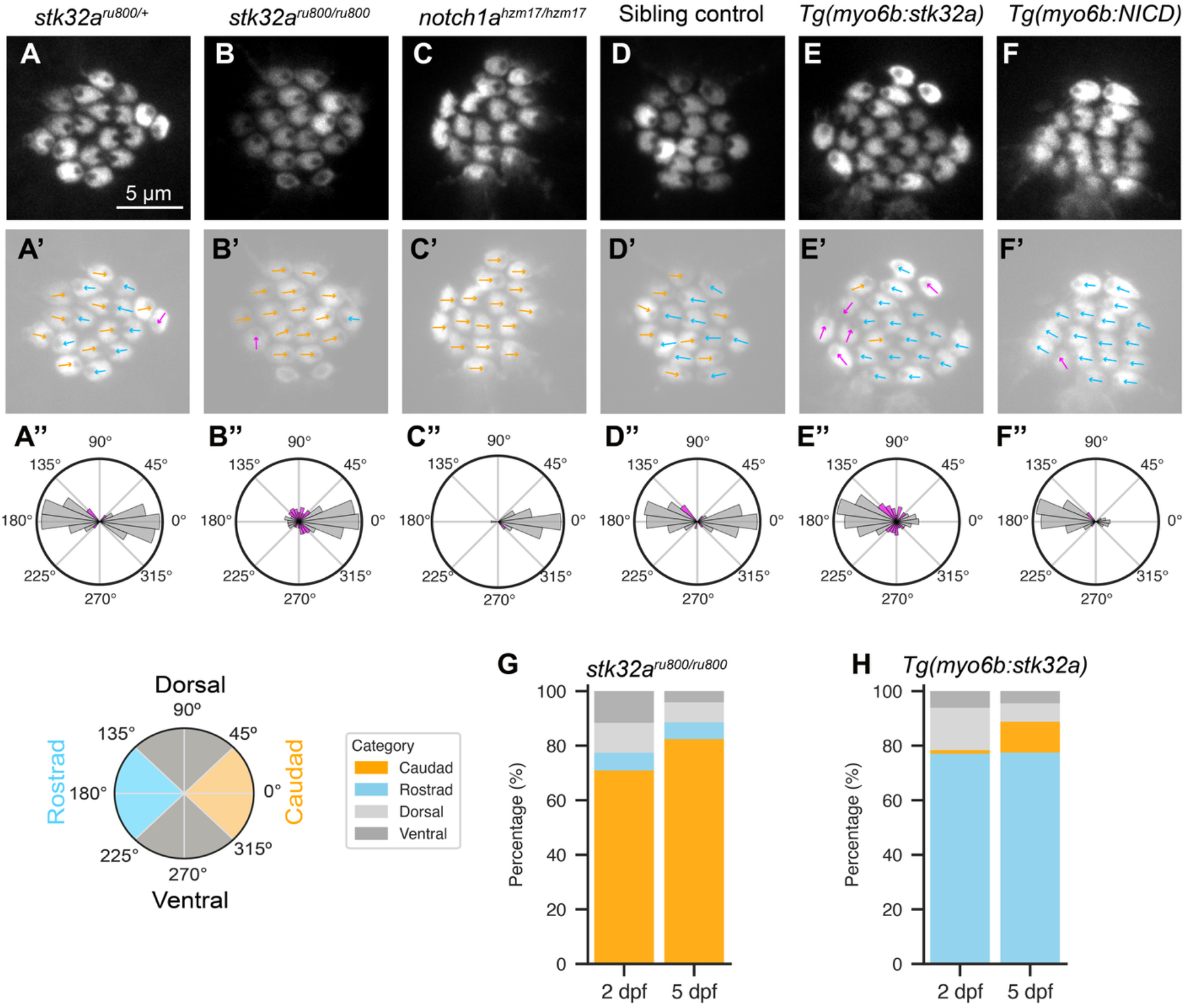
Hair bundle polarization at 5 dpf, related to Figure 4. **(A-F)** Images of the apical surface of hair cells in the specified genetic backgrounds at 5 dpf; a single optical section or a max Z projection of 2-3 slices is shown for ease of visualization of hair bundle polarization. Scale bar, 5 µm. **(A’-F’)** The same images as above, dimmed and overlaid with arrows representing the polarization of the hair bundles: orange: caudad, blue: rostrad, pink: off-axis (dorsal or ventral-polarized). **(A’’-F’’)** Rose plots depicting polarity spreads at 5dpf, generated as in Figure 4; magenta bar color indicates off-axis polarization. (A) *stk32a*^*ru800/+*^ siblings (n = 294 hair cells from 19 neuromasts in 5 larvae). (B) *stk32a*^*ru800/ru800*^ mutants (n = 296 hair cells from 19 neuromasts in 5 larvae). (C) *notch1a*^*hzm17/hzm17*^ mutants (n = 167 hair cells from 11 neuromasts in 4 larvae). (D) Sibling non-expressing controls (n = 320 hair cells from 18 neuromasts in 5 larvae). (E) *Tg(myo6b:stk32a)* (n = 311 hair cells from 18 neuromasts in 5 larvae). (F) *Tg(myo6b:NICD)* (n = 283 hair cells from 19 neuromasts in 5 larvae). **(G-H)** Bar plots showing the proportion of caudad, rostrad, dorsal, or ventral-polarized hair bundles for the given genotype at 2 and 5 dpf. Note that 2 dpf data are displayed in Figure 4. A Fisher’s Exact test (with Holm-correction) was performed on collapsed data: caudad/rostrad (on axis) vs. dorsal/ventral (off axis). *stk32a(ru800/ru800)* mutants showed a significant decrease in off-axis polarity from 2 to 5 dpf (**, p < 0.01), as did *Tg(myo6b:stk32a)* neuromasts (**, p < 0.01).

**Supplemental Figure S8.**
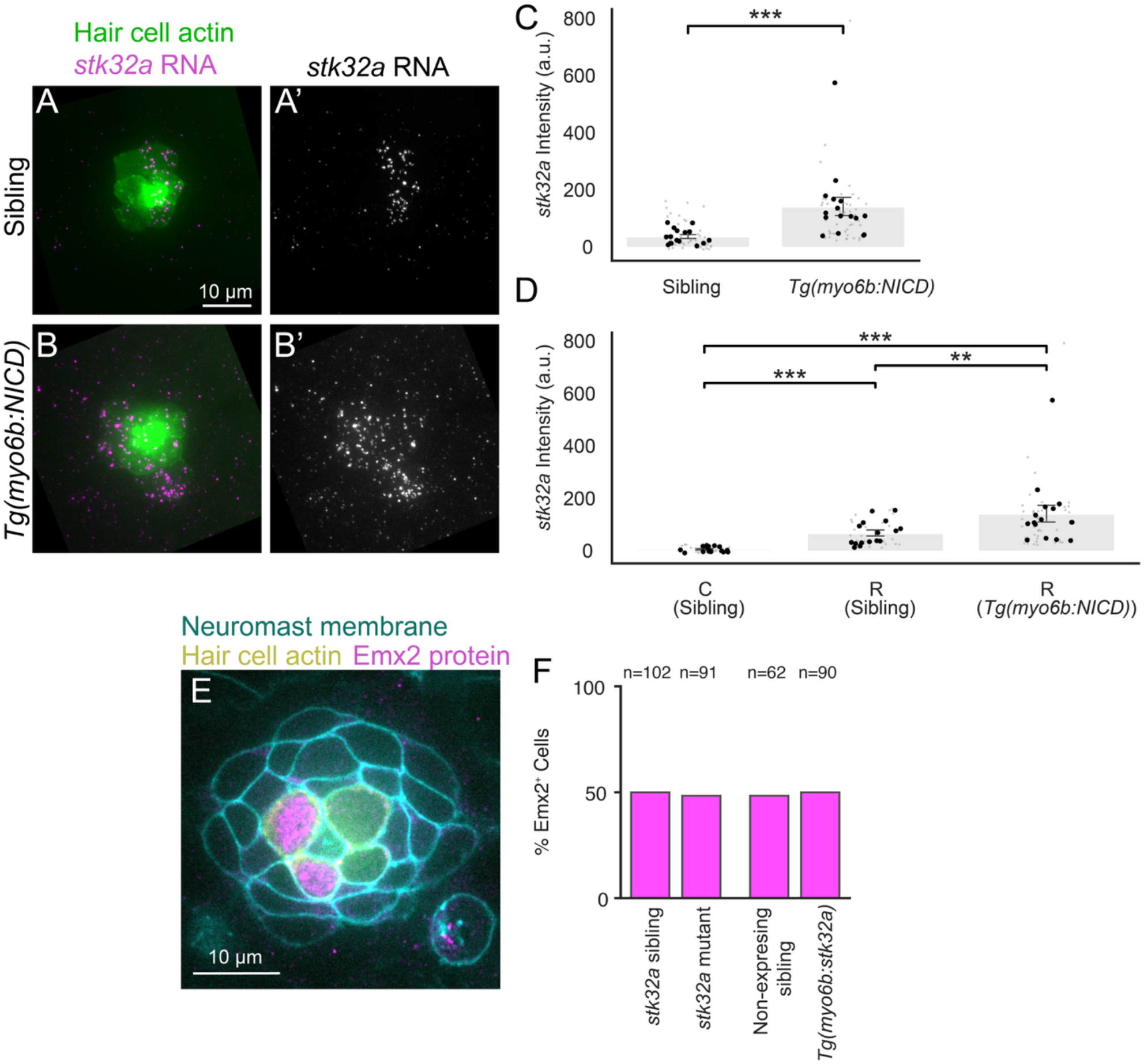
Notch-mediated regulation of *stk32a*, related to Figure 4. **(A, B)** Example larvae stained by hybridization chain reaction (HCR) *in situ* with probes against *stk32a*, shown in a sibling larva (A) and a *Tg(myo6b:NICD)* larva (B). Images are maximum intensity projections through planes containing hair cells, rotated to the align with the organ’s axis. *stk32a* RNA (magenta) is overlaid with the *Tg(myo6b:actb1-EGFP)* transgene (green) marking hair cell actin. **(A’, B’)** *stk32a* RNA expression alone (gray) from the same images. All images were intensity-adjusted individually for each channel to visualize presence of *stk32a* rather than relative intensity of staining. Scale bar, 10 µm. **(C)** *stk32a* expression in mature hair cells. Intensity values were measured in individual mature hair cells (maximum intensity z-projection) in raw unprocessed images; a local background measurement was taken and subtracted separately for each neuromast image. Bars represent mean ± SEM of neuromast means. Gray points show individual cells; black points show neuromast means. n=16 neuromasts (52 cells) for siblings, n=16 neuromasts (52 cells) in *Tg(myo6b:NICD)* larvae, ***p < 0.001, Mann-Whitney U test. **(D)** *stk32a* expression in mature hair cells divided by polarity. Siblings have both caudad- and rostrad-polarized hair cells, whereas *Tg(myo6b:NICD)* have just rostrad-polarized cells. Bars represent mean ± SEM of neuromast means. Gray points show individual cell intensities; black points show neuromast means. n=16 neuromasts (26 cells) for C (Sibling), n=16 neuromasts (26 cells) for R (Sibling) and n=16 neuromasts (52 cells) for *Tg(myo6b:NICD)*. **p < 0.01, ***p < 0.001, Mann-Whitney U test with Holm correction for multiple comparisons. **(E)** Emx2 protein localization (magenta) in a single optical section of a *stk32a*^*ru800/ru800*^ neuromast. Cell membranes are labeled with *Tg(cldnb:lyn-mScarlet)* in cyan; hair cell actin labeled with *Tg(myo6b:actb1-EGFP)* in yellow. Scale bar, 10 µm. **(F)** Percentage of Emx2-positive hair cells for each genotype. Numbers above bars indicated total hair cells analyzed. Data from same experiments as Figure 4G-J.

**Supplemental Figure S9.**
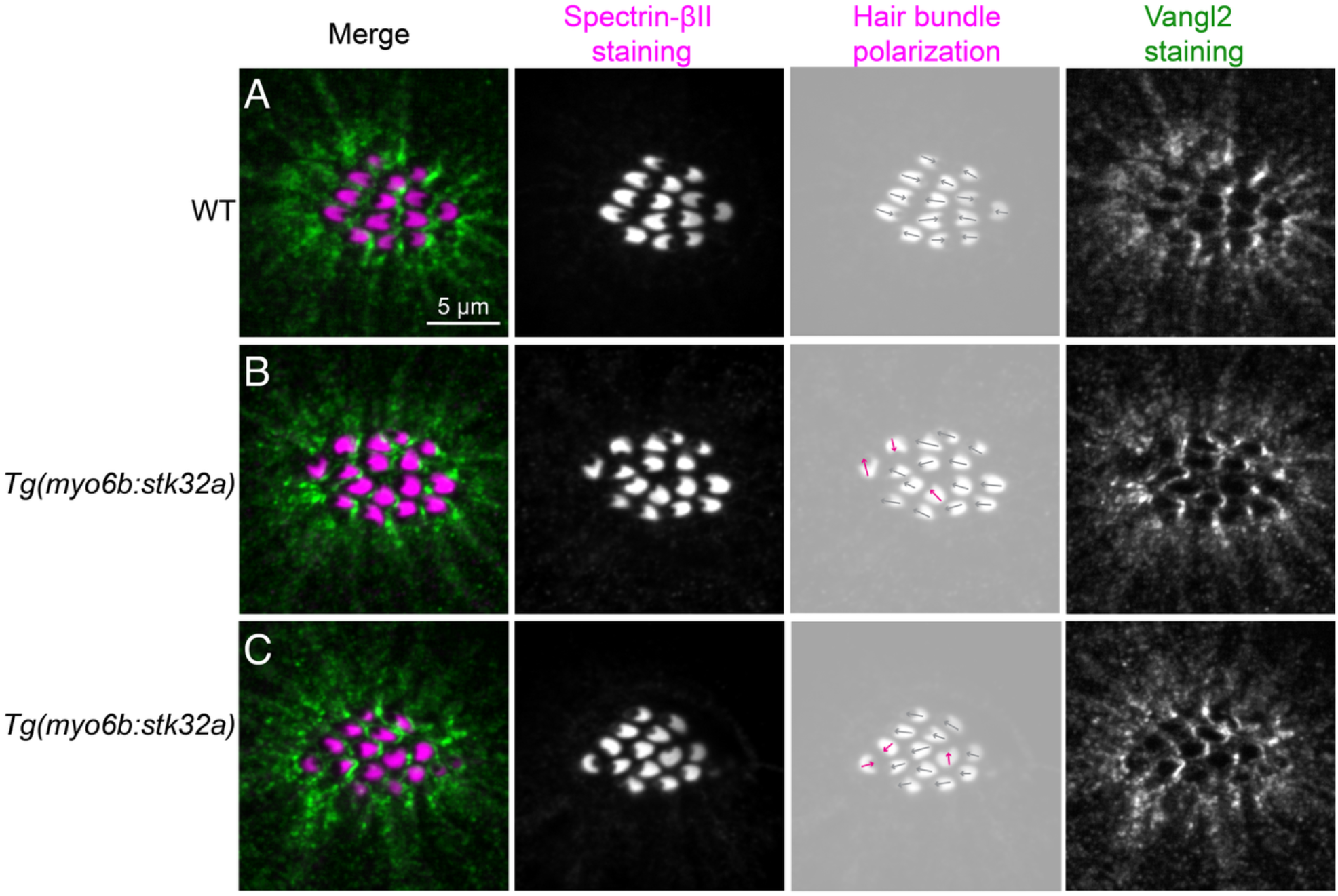
Vangl2 localization in *Tg(myo6b:stk32a)* neuromasts, related to Figure 4. Vangl2 protein localization in *Tg(myo6b:stk32a)* neuromasts. **(A)** Representative control neuromast (sibling of transgenic larva). Images were contrast-adjusted and rotated to align with the organ’s axis. Merged image is shown as a maximum-intensity projection of nine 0.2-μm z-slices encompassing the apical surface showing Spectrin-βII (magenta), which labels actin-rich hair bundles but is excluded from the kinocilium, enabling assessment of bundle polarity, and Vangl2 (green). <u>Middle panels</u>: A single optical section of Spectrin-βII is shown in grayscale to facilitate visualization of hair bundle polarization. A dimmed image in grayscale with arrows indicating hair-bundle polarization; gray indicates rostrad-polarized hair bundles and magenta arrows indicate either off-axis or caudad-polarized hair bundle polarities. <u>Right panel</u>: Vangl2 shown in grayscale (maximum-intensity projection of 9 slices, as in merged image). Images were intensity-adjusted independently for each sample to facilitate visualization of relative intensity patterns within a sample. Scale bar, 5 μm. **(B-C)** Same as in A, but for two representative *Tg(myo6b:stk32a)* neuromasts.

**Supplemental Figure S10.**
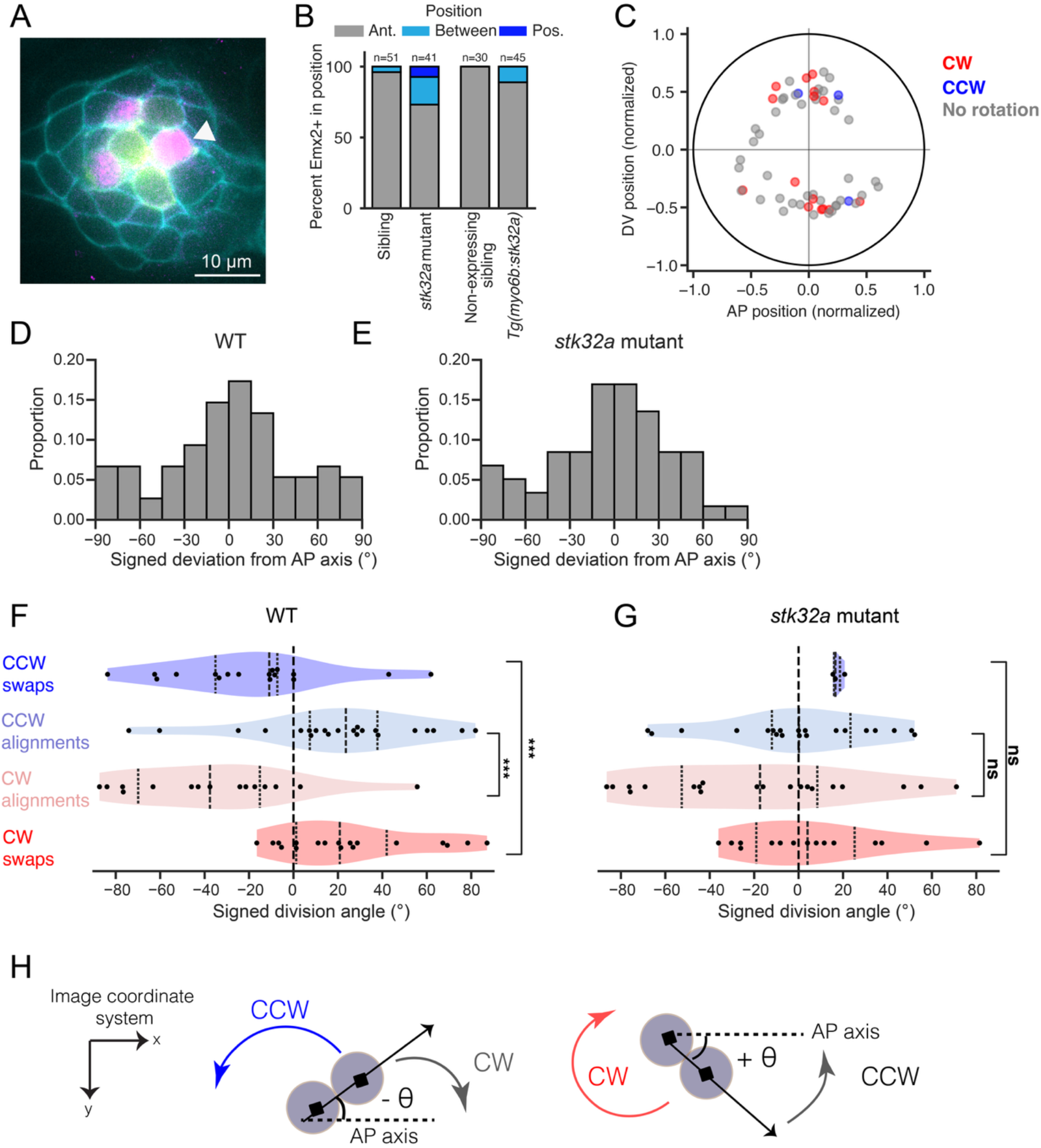
Extended data from Figure 5. **(A)** A single optical section of a 2 dpf neuromast showing Emx2 protein localization by antibody staining (magenta) in a *stk32a*^*ru800/ru800*^ mutant neuromast with a mispositioned Emx2-positive hair cell (white arrowhead). Cell membranes are labeled with *Tg(cldnb:lyn-mScarlet)* marked in cyan, and hair cell actin with *Tg(myo6b:actb1-EGFP)*, marked in yellow. Scale bar, 10 µm. **(B)** Percent of Emx2+ hair cells within each neuromast categorized by their position relative to all other hair cells in that neuromast: anterior (gray), between (light blue), or posterior (dark blue). Numbers above bars indicate the total Emx2+ cells analyzed; data are from the same cells as in figure 4G-J, except neuromasts with an odd number of hair cells were excluded. A two-sided Fisher’s Exact test was used to compare the proportion of Emx2+ cells located on the anterior versus not on the anterior of all other cells, comparing mutants and their siblings (**p < 0.01), and *Tg(myo6b:stk32a)* neuromasts and their siblings (ns, p ≥ 0.05), Holm-corrected. **(C)** Division locations of *stk32a*-mutant progenitors (normalized to neuromast radius). Color indicates swap outcome: red, clockwise swap; blue, counter-clockwise swap; gray, non-swapping pair. **(D-E)** Histogram depicting the spread of division angles for WT and *stk32a*-mutant pairs folded to [-90º,90º] in WT hair cell pairs (D) or *stk32a* mutant pairs (E). **(F-G)** Violin plot showing division angles of pairs classified as swaps vs. non-swaps (alignments) classified by direction of rotation CW or CCW in WT **(F)** or *stk32a* mutant **(G)** pairs. Cell pairs rotating ≥ 360º were excluded (3/75 for WT and 1/59 for *stk32a* mutant). Individual cell pair data are overlaid (one point per cell pair). Statistical comparisons between CW and CCW were performed separately for swaps and alignments using two-sided Mann–Whitney U tests with Holm correction (two tests per genotype); ***p < 0.001, ns p ≥ 0.05. **(H)** Expected rotation direction given division sign to minimize movement; negative angle: swap CCW, align CW; positive angle: swap CW, align CCW. This preference is evident in WT pairs but not in cell pairs from *stk32a* mutants.

## Supplementary Information

**Supplemental Movie 1: 3D rendering of full neuromast segmentation**. Movie of cell-pair rotation shown in Figures 1C and 1D, showing both XY and XZ views with the full neuromast rendered in gray and the tracked cell pair colored in blue and orange.

**Supplemental Movie 2: 3D rendering of cell-pair rotation, zoomed in on swapping cell pair**. Zoomed view of the same cell pair shown in Supplemental Movie 1, corresponding to Figures 1C and 1D, showing both XY and XZ views.

**Supplemental Movie 3: 3D rendering of cell-pair that swaps**. Related to Figure 1F. Movie of the cell pair whose angular trajectory is shown in Figure 1F.

**Supplemental Movie 4: 3D rendering of cell-pair that does not swap**. Related to Figure 1G. Movie of the cell pair whose angular trajectory is shown in Figure 1G.

**Supplemental Movie 5: 3D rendering of cell-pair that aligns and swaps**. Related to Figure 1H. Movie of the cell pair whose angular trajectory is shown in Figure 1H.

**Supplemental Table 1:** Differential gene expression between *notch1a* mutants and siblings/wild type (all hair cell data). Positive log_2_ fold change indicates upregulation in *notch1a* mutants. Results are filtered for significant changes (see Methods).

**Supplemental Table 2:** Differential gene expression between *notch1a* mutants and siblings/wild type (young hair cell data). Positive log_2_ fold change indicates upregulation in *notch1a* mutants. Results are filtered for significant changes (see Methods).

**Supplemental Table 3:** Differential gene expression analysis between Notch-OFF and Notch-ON trajectories using tradeSeq *patternTest* on young hair cell pseudotime trajectories. Gene list sorted by adjusted p-value and median fold change (fcMedian).

**Supplemental Table 4:** F0 CRISPR screen summary.

## Notes

### Competing Interest Statement

The authors have declared no competing interest.

### Summary of Updates

Physical model added (Figure 6); title slightly changed; data availability section added; some copy editing.

